# Syntactic Processing Engages the Semantic Control Network

**DOI:** 10.1101/2025.10.31.685882

**Authors:** Qianwen Chang, Elizabeth Jefferies, Rebecca L. Jackson

**Author notes:** These authors contributed equally (Elizabeth Jefferies, Rebecca L. Jackson).

## Abstract

The relationship between semantics and syntax is highly contested. Neuroimaging evidence has offered conflicting views on whether these domains are neurally separable, in part because prior work has not distinguished two key components of semantic cognition: *semantic representation* and *semantic control*. In this study, we conducted an activation likelihood estimation (ALE) meta-analysis to identify regions consistently engaged by more versus less demanding syntactic processing, and compared these directly with regions recruited by demanding versus less demanding semantic processing. Demanding syntactic processing engaged regions across the semantic control network (SCN), including the inferior frontal gyrus (IFG) and insula (particularly on the left), left posterior temporal cortex (posterior superior temporal sulcus and middle temporal gyrus), and bilateral dorsomedial prefrontal cortex. No regions outside the SCN responded significantly more to syntactic demands than semantic control. Within the SCN, there was significantly less reliable activation for syntactic than semantic control demands in some anterior (pars triangularis) and ventral (pars orbitalis) portions of the left IFG and a region in posterior fusiform and inferior temporal gyrus. Greater involvement for syntax than semantic control was identified in a small region of posterior left IFG (pars opercularis) within the SCN. Moreover, both controlled semantic and syntactic processing showed multimodal responses to auditory and visual stimuli in the left IFG and posterior temporal cortex. Together, these findings suggest that syntactic processing is distinct from semantic representation, yet demanding syntax engages portions of the SCN, reflecting a shared need to access and manipulate stored semantic knowledge for flexible, context-sensitive use.

## 1 Introduction

The relationship between syntax and semantics has long been debated. Syntactic processing involves analysing the relationship between words, including sentential structure (Friederici & Kotz, 2003), while semantic cognition allows comprehension of their meaning. Both semantics and syntax rely on statistical learning (Saffran, 2020; Thompson & and Newport, 2007), and can be learnt simultaneously (Poletiek et al., 2021). Despite this, the relationship between these processes remains controversial. It is argued that syntax and semantics are orthogonal (Chomsky, 1957), that is, it is possible to manipulate either the syntactic structure or the meaning of a sentence independently without affecting the other. For example, *The dog [the cat chases] runs* and *The cat [the dog chases] runs* have identical syntactic structure but different meanings (Poletiek et al., 2021). Indeed, semantic dementia patients show a loss of semantic concepts with preserved syntax (Landin-Romero et al., 2016; Rochon et al., 2004), providing strong evidence that syntax and semantics are independent. In addition, syntax appears language-specific, while the semantic network responds similarly to both verbal and nonverbal stimuli, including pictures and actions (Canini et al., 2016; Krieger-Redwood et al., 2015; Schnur et al., 2009; Sitnikova et al., 2014) and damage results in multimodal semantic deficits (Corbett et al., 2009; Jefferies & Lambon Ralph, 2006). However, simply learning to predict the next word in a sentence provides large language models with remarkable semantic and syntactic abilities, without the need to rely on distinct processes or regions (Arana et al., 2024; Contreras Kallens et al., 2023). Due to the compelling evidence on both sides of this debate, the relationship between semantic and syntactic processing remains unclear (Friederici & Weissenborn, 2007; Kako & Wagner, 2001; Pylkkänen, 2019).

Recent research may transform this debate by highlighting the multifactorial nature of semantic cognition. Semantic processing relies on the interaction between two separable processes, semantic representation and semantic control (Lambon Ralph et al., 2017). The semantic representation system encodes generalisable semantic information across contexts, while the semantic control system manipulates activation within the representation system in specific contexts, focusing on task-relevant knowledge and inhibiting task-irrelevant knowledge (Chiou et al., 2018; Hoffman et al., 2018; Jefferies, 2013; Lambon Ralph et al., 2017). This semantic control system is at least somewhat separable from domain-general executive control, relying upon regions preferentially activated for the controlled access and manipulation of meaningful over meaningless stimuli, even when similar task processes are performed (Hodgson et al., 2024). Semantic dementia patients have focal damage to bilateral anterior temporal lobes, resulting in a specific impairment of multimodal semantic representation, rather than semantic control (Jefferies et al., 2008; Jefferies & Lambon Ralph, 2006; Lambon Ralph & Patterson, 2008; Patterson et al., 2007). Therefore, their preserved syntax only demonstrates that syntax is dissociable from semantic representation. Damage to left inferior prefrontal or posterior temporal regions results in impairment of semantic control, referred to as semantic aphasia (Jefferies & Lambon Ralph, 2006; Thompson et al., 2022). However, the relationship between this impairment and syntactic abilities has not been systematically investigated. Therefore, the relationship between semantic control and syntax remains unclear.

Neuroimaging studies have shown that syntactic and semantic processing recruit overlapping frontal-temporal networks, including the left inferior frontal gyrus (IFG) and posterior temporal cortex (Bautista & Wilson, 2016; Fedorenko et al., 2020; Friederici, 2012; Heim et al., 2008). Syntax is primarily associated with left IFG, posterior superior and middle temporal gyrus (pSTG/MTG), and inferior parietal lobe (IPL) (Friederici et al., 2003; Friederici & Kotz, 2003; Hagoort & Indefrey, 2014; Rodd et al., 2015; Turker et al., 2023; Tyler et al., 2011), while semantic control recruits bilateral IFG (with a preference for the left), left posterior middle and inferior temporal gyrus (pMTG/ITG), and bilateral dorsomedial prefrontal cortex (dmPFC) (Gao et al., 2021; Jackson, 2021; Jefferies, 2013; Noonan et al., 2013; Wang et al., 2018). There may therefore be overlap between these neural networks, in frontal and temporal cortices, which could reflect shared control processes supporting demanding syntactic and semantic processing. Although the two processes appear to recruit similar regions, dissociations have been reported in the left IPL and IFG. The left IPL is strongly associated with syntactic processing (Lee & Newman, 2010; Tyler et al., 2011), but responds less reliably to semantic control (Jackson, 2021). In addition, prior work has demonstrated relative dissociations between anterior left IFG for semantic processing and posterior IFG for syntactic processing (Friederici, 2011; Goucha & Friederici, 2015; Schell et al., 2017; Vigneau et al., 2006), although these studies did not consider semantic and syntactic control demands.

Formal meta-analyses are needed to systematically identify the relationship between semantic control and syntactic processing, since existing comparisons have not considered subcomponents of semantic cognition (Turker et al., 2023). One exception is Rodd et al. (2015) who investigated semantic and syntactic processing across visual and auditory modalities, contrasting high-demands with easier tasks. However, the studies available at the time left these comparisons underpowered (26 semantic and 28 syntactic studies), and they were unable to compare across the two modalities.

In the present study, we aimed to identify the neural correlates of syntactic processing and semantic control to examine two key questions: Firstly, to what extent does demanding syntactic processing recruit regions of the semantic control network versus additional domain-general or syntax-specific areas? Secondly, are the regions engaged by syntactic processing multimodal, responding to both auditory and visual stimuli, and how does this compare to the effects of modality in semantic control regions? To address these questions, we focused on conditions of varying demand in both semantic and syntactic domains, following a novel meta-analytic approach inspired by Rodd et al. (2015). The syntactic contrast was constructed in a manner analogous to standard semantic control contrasts (e.g., Jackson, 2021; Noonan et al., 2013), allowing direct comparison of controlled processing across the two domains. Previous studies have reported multimodal responses in left IFG and posterior temporal cortex for both semantic control and syntactic tasks (Buchweitz et al., 2009; Corbett et al., 2009; Jackson, 2021; Jefferies & Lambon Ralph, 2006), yet the precise relationship between multimodal semantic and syntactic regions remains unclear.

## 2 Method

This meta-analysis was performed and reported following the Preferred Reporting Items for Systematic Reviews and Meta-Analyses (PRISMA) statement (Moher et al., 2009; Page et al., 2021). The study screening and selection process was performed in Covidence (www.covidence.org), a systematic review software.

### 2.1 Inclusion and exclusion criteria

#### 2.1.2 Syntactic processing

For syntactic processing, studies were taken from two sources: previous meta-analyses and a new literature search. First, studies identified from previous meta-analyses in syntactic processing (Hagoort & Indefrey, 2014; Heard & Lee, 2020; Rodd et al., 2015; Vigneau et al., 2006, 2011; Zaccarella et al., 2017) were included. While some of these meta-analyses focused on specific operations in syntactic processing (e.g., merge in Zaccarella et al., 2017), others took a broader view. For example, Hagoort and Indefrey (2014) examined semantic and syntactic processing at the sentence level, categorising syntactic manipulations into complexity, violation, and ambiguity. Similarly, Rodd et al. (2015) compared semantic and syntactic processing, encompassing comprehensive syntactic manipulations (i.e., violation, complexity, ambiguity, and cross-modal priming). They used stringent inclusion criteria — excluding studies where contrasts included both syntactic and semantic processing (e.g., sentences vs. word lists, and sentences vs. ‘jabberwocky’ sentences) — to avoid potential confounds between these two domains. These previous meta-analyses covered the literature until 2014. To extend this, we therefore conducted a literature search in Web of Science from 2014 to May 2024. To identify as much related literature as possible, we used the search string: (fMRI OR functional magnetic resonance imaging OR PET OR positron emission tomography) AND (syntax OR syntactic OR grammar OR grammatical OR sentence). After removing duplicates from the two sources, 1,400 studies were included in the initial screening.

The initial screening was performed based on titles and abstracts. Studies that did not meet the following criteria were excluded in this initial screening: (1) peer-reviewed journal articles written in English; (2) empirical studies (not reviews or meta-analyses); (3) task-based fMRI or PET studies; (4) studies focused on healthy young adults (aged 18-50 years), including healthy control groups in patient studies; (5) studies using tasks where syntactic processing was the major cognitive process. After the initial screening stage, 255 studies passed to the second screening stage.

The second screening stage was performed based on the full texts. Studies had to meet the criteria described above, as well as the following criteria: (1) whole-brain coverage; (2) univariate activation analysis without small volume correction; (3) peak coordinates reported in Talairach or Montreal Neurological Institute (MNI) stereotaxic space; (4) stimuli were sentences, word lists or word pairs in visual or auditory modalities; (5) contrasts were more demanding > less demanding syntactic processing, including ambiguous > unambiguous, complex > simple sentence structures and syntactically incorrect > correct. Studies using low-level references (e.g., fixation, rest, or other non-linguistic stimuli) were excluded, as these involve cognitive processes other than language processing. Moreover, following Rodd et al. (2015), studies where the contrasts resulted in differences in both semantic and syntactic processing (e.g., sentences vs. word lists, and sentences vs. ‘jabberwocky’ sentences) were excluded, to promote the separation of these domains. Thus, all included studies involved an active syntactic task performed on verbal stimuli, and contrasted a more demanding over a less demanding condition. This allowed us to isolate brain regions engaged by syntactic demands. Cross-modal priming studies were not included in the present study. Following Hagoort and Indefrey (2014), studies were further classified into 3 categories: syntactic violation (e.g., subject-verb disagreement > agreement), ambiguity (e.g., word-class ambiguity), and complexity paradigms (e.g., passive > active sentences, non-canonical > canonical word-order, center-embedding > left/right-branching, object-relative > subject-relative clauses) based on syntactic manipulations. Across these studies, participants performed various tasks to probe syntactic processing, such as passive listening/reading, grammaticality judgment, comprehension/verification, and picture-sentence matching tasks. Multiple contrasts reported for the same participant sample were analysed as a single contrast following the recommendation of Müller et al. (2018).

A total of 66 experiments with 513 foci were included in the formal meta-analysis (Figure 1). In follow-up analyses, these were divided to compare the 21 experiments (108 foci) in the auditory modality with the 43 experiments (371 foci) in the visual modality, and two experiments (34 foci) involving both modalities. All 66 experiments were included for the analysis of general syntactic processing, while the two experiments involving both modalities were excluded from modality-specific analyses. Additionally, following Hagoort and Indefrey (2014), the particular syntactic manipulation was compared by contrasting the 50 experiments (346 foci) on complexity and 14 experiments (151 foci) on violation. The two experiments (16 foci) in ambiguity were not assessed independently, as there was an insufficient number of studies for a meta-analysis (as we utilised stricter inclusion criteria to disambiguate syntax and semantics than prior investigation). All experiments included are described, and the peaks are provided in Supplementary Table 1.

**Figure 1.**
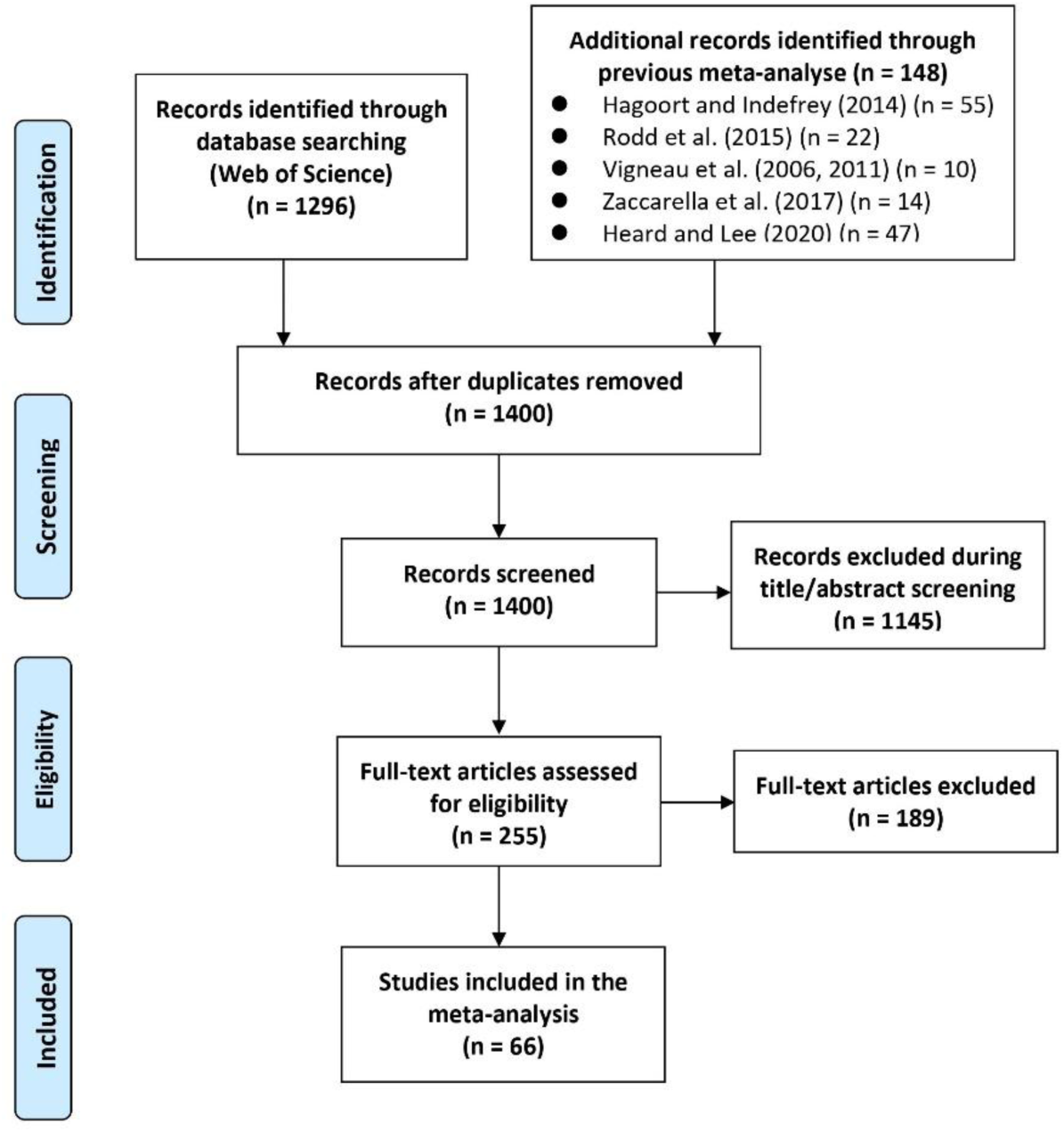
PRISMA flow chart for syntactic processing

#### 2.1.1 Semantic control

The regions consistently activated for syntactic demands were compared to those implicated in semantic control. For semantic control, we sourced the same studies as a recent meta-analysis on semantic control (Jackson, 2021; see also Hodgson et al., 2023). This meta-analysis contrasted more over less demanding semantic processing across both verbal and nonverbal stimuli in visual and auditory modalities. However, to focus on the language domain, we restricted our analysis to studies using verbal stimuli in both visual and auditory modalities (7 experiments with 46 foci using non-verbal stimuli in visual modality were excluded). This resulted in a total of 82 experiments with 871 foci, including 65 experiments (712 foci) in the visual modality and 17 experiments (159 foci) in the auditory modality.

### 2.2 Activation likelihood estimation

The meta-analyses were performed in GingerALE 3.0.2 (https://brainmap.org) using the activation likelihood estimation method (ALE; Eickhoff et al., 2009, 2012, 2017; Turkeltaub et al., 2012). Talairach foci were first converted to MNI standard space using the tal2icbm_spm transformation implemented in GingerALE. All analyses were performed in MNI152 space. The ALE method identifies regions of consistent activation across studies by modelling reported foci as Gaussian probability distributions. First, a model activation map was generated for each experiment, where each activation focus was modelled as a three-dimensional Gaussian probability distribution centred at its reported coordinates. To account for spatial uncertainty, the Gaussian distribution’s full-width at half-maximum (FWHM) was adjusted according to each study’s sample size (larger sample sizes resulting in smaller FWHM values, indicating greater spatial precision). Next, an ALE map was generated by computing the union of model activation maps across all experiments, resulting in an ALE score for each voxel. This map represents the convergence of activation probabilities across studies, with each voxel’s ALE score indicating the likelihood of consistent activation at that location.

Independent ALE analyses were performed to identify the areas consistently involved in syntax and semantic control. ALE scores were thresholded at *p* < 0.001 at the voxel-level and FWE corrected *p* < 0.05 at the cluster-level, with 10,000 permutations (Müller et al., 2018). Contrast and conjunction analyses were conducted on the resulting maps to reveal the distinct and shared neural underpinnings of semantic control and syntactic processing. The conjunction analysis simply identifies any voxels present in both thresholded maps. The contrast analysis identifies regions more likely to activate in one condition over the other by subtracting one thresholded map from the other and assessing whether the resulting differences are significantly larger than would be expected by chance. Contrast scores were thresholded at voxel-level *p* < 0.001 with 10,000 permutations, cluster volume > 20 mm^3^. The two syntactic manipulations (syntactic complexity and violation) were then analysed separately and compared. Additionally, we examined the pattern of activation for syntax and semantic control in the visual and auditory modalities, and compared these modality-specific patterns (visual vs. auditory syntax and visual vs. auditory semantic control).

## 3 Results

### 3.1 Syntactic processing regions

The regions consistently activated during syntactic processing are shown in Figure 2. Peak coordinates are reported in Table 1. The largest cluster is in the left IFG, including pars opercularis and pars triangularis, extending to the insula and precentral gyrus. The strongest activation likelihood is in pars opercularis. A second cluster is in the left posterior middle and superior temporal gyri, focused on the posterior superior temporal sulcus (pSTS). Additional activations are in the left IPL (including angular gyrus and supramarginal gyrus) and bilateral dmPFC, including supplementary motor area (SMA). Two clusters are identified in the right hemisphere, in the insula and IFG, including pars opercularis and pars triangularis. This result reveals a distributed, left-dominant network for syntactic processing, focused around IFG, dmPFC and dorsal posterior temporal cortex. Analyses comparing the type of syntactic manipulations are presented in the Supplementary Materials (Supplementary Figure 1 and Supplementary Table 2) only, as most studies included variation in syntactic complexity, rather than violation or ambiguity. Whether different syntactic assessments would result in changes to this network should be explored in further work.

**Figure 2.**
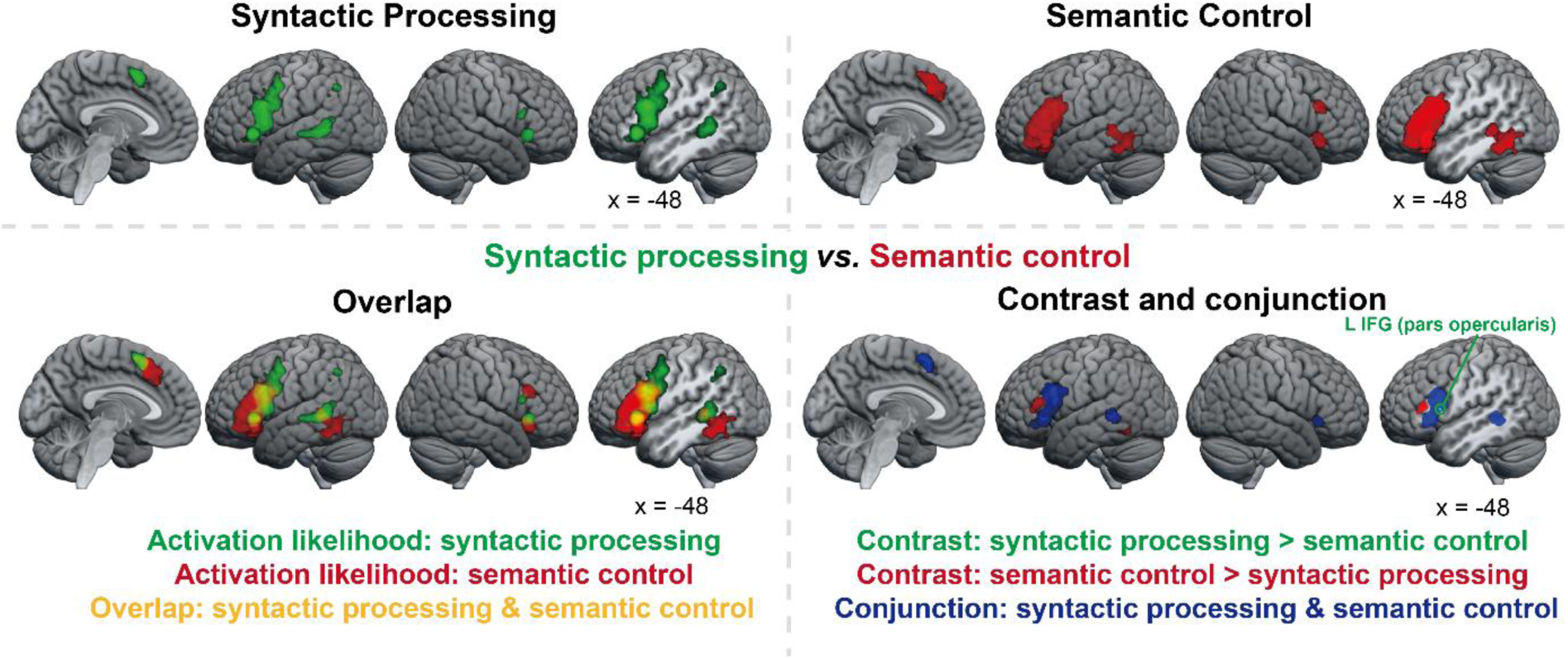
Top: Activation likelihood estimation maps for syntactic processing (top left, shown in green) and semantic control (top right, shown in red) at a voxel-level *p* < 0.001, cluster-level FWE corrected *p* < 0.05, with 10,000 permutations. Bottom: Overlap of ALE results of syntactic processing and semantic control (bottom left, overlap in yellow). Contrast and conjunction analysis between syntactic processing and semantic control at a voxel-level *p* < 0.001 with 10,000 permutations, cluster volume > 20 mm^3^ (bottom right, syntactic processing > semantic control in green, semantic control > syntactic processing in red, conjunction in blue).

**Table 1.**
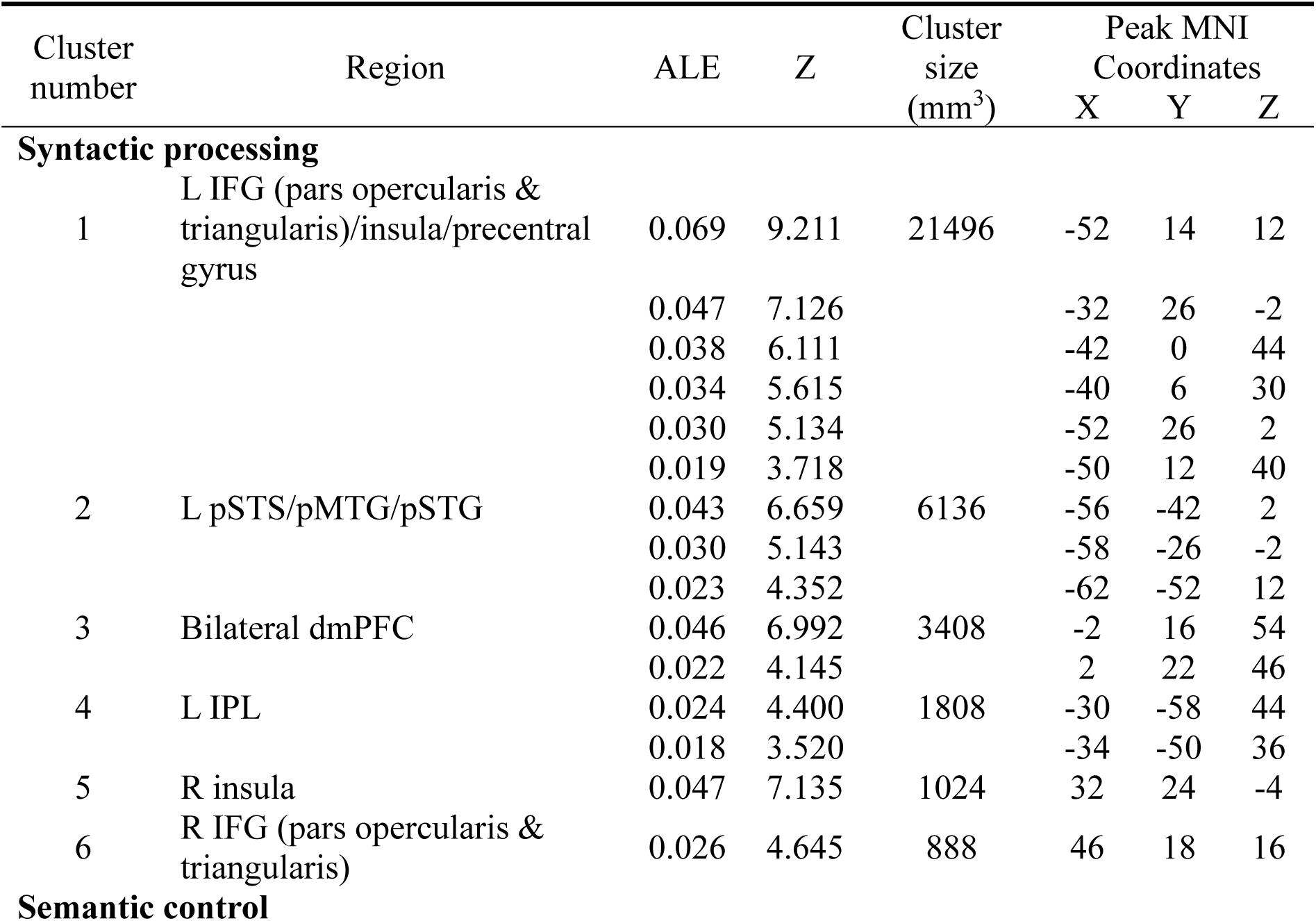
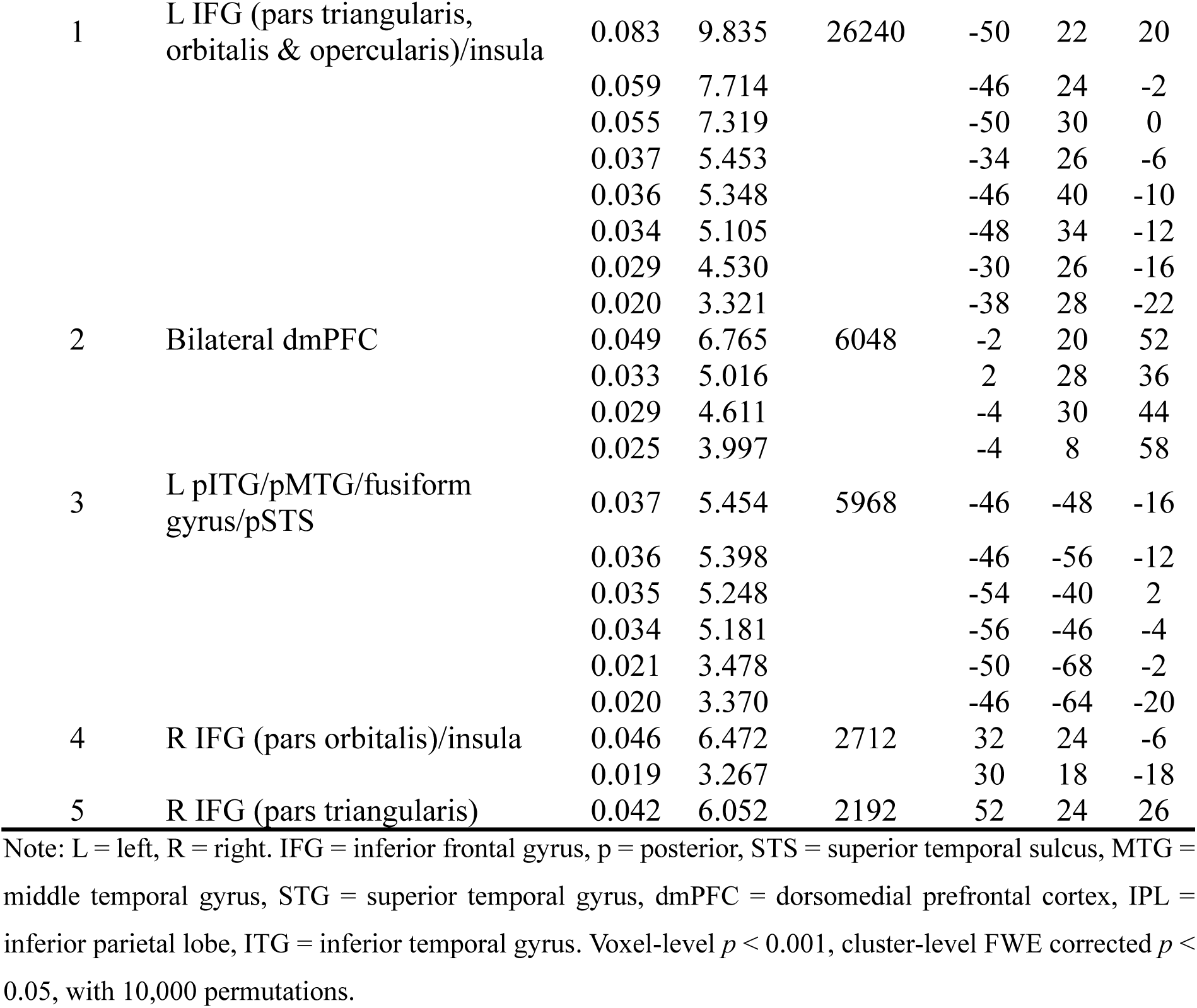
Activation likelihood estimation values for syntactic processing and semantic control.

### 3.2 Comparing syntactic processing regions to semantic control

Semantic control also relies upon a distributed left-dominant network with crucial nodes in the left prefrontal and posterior temporal cortices. To directly compare the regions implicated in semantic and syntactic demands, we replicated a recent semantic control meta-analysis, using only the verbal stimuli (Figure 2 and Table 1). As found previously (Jackson, 2021; Noonan et al., 2013), bilateral IFG, insula, and dmPFC are consistently implicated in semantic control, alongside a left posterior temporal cortex region, covering the lateral surface from the posterior STS, through MTG and ITG into fusiform gyrus.

Directly comparing the regions consistently implicated in the two domains highlighted substantial overlap in left IFG extending to the insula and precentral gyrus, left dmPFC, left pMTG/pSTS, and right insula (see Figure 2 and Table 2). Formal contrast analyses nevertheless identified some areas of difference. Compared to syntactic processing, semantic control showed greater activation in left dorsal anterior IFG (pars triangularis), as well as posterior fusiform and inferior temporal gyri, and a small cluster in ventral IFG (pars orbitalis). In contrast, syntactic processing showed greater activation in a small region of left IFG (pars opercularis), overlapping the area consistently implicated in both processes.

**Table 2.** Contrast and conjunction analyses between syntactic processing and semantic control.

| Cluster number | Region | Cluster size (mm<sup>3</sup>) | Peak MNI Coordinates |  |  |
| --- | --- | --- | --- | --- | --- |
|  |  |  | X | Y | Z |
| **Syntactic processing vs. Semantic control** |  |  |  |  |  |
| *Contrast analysis: syntactic processing > semantic control* |  |  |  |  |  |
| 1 | L IFG (pars opercularis) | 32 | -54 | 16 | 10 |
| *Contrast analysis: semantic control > syntactic processing* |  |  |  |  |  |
| 1 | LIFG (pars triangularis) | 1944 | -49 | 32 | 13 |
| 2 | L fusiform gyrus/ITG | 352 | -48 | -58 | -16 |
|  |  |  | -52 | -60 | -10 |
|  |  |  | -46 | -50 | -16 |
| 3 | L IFG (pars orbitalis) | 24 | -28 | 32 | -18 |
| *Conjunction: syntactic processing & semantic control* |  |  |  |  |  |
| 1 | L IFG (pars opercularis & triangularis)/insula/precentral gyrus | 12328 | -52 | 18 | 18 |
|  |  |  | -52 | 26 | 0 |
|  |  |  | -34 | 26 | -4 |
|  |  |  | -42 | 6 | 26 |
| 2 | L dmPFC | 2032 | -2 | 16 | 54 |
| 3 | L pMTG/pSTS | 1680 | -54 | -42 | 2 |
|  |  |  | -56 | -46 | -2 |
| 4 | R insula | 912 | 32 | 24 | -6 |

Note: L = left, R = right. IFG = inferior frontal gyrus, ITG = inferior temporal gyrus, dmPFC = dorsomedial prefrontal cortex, p = posterior, MTG = middle temporal gyrus, STS = superior temporal sulcus. Voxel-level $p < 0.001$ with 10,000 permutations, cluster volume $> 20 \text{ mm}^3$ .

### 3.3 Comparing syntactic processing regions to semantic representation and multiple-demand network

Syntactic demands activate the semantic control network, yet this may not be the full picture. How can we explain the differences identified between syntactic and semantic demands, albeit in the context of large overlap? Does demanding syntactic processing recruit semantic regions more generally outside of the SCN, and/or does it rely on additional domain-general cognitive control processes beyond semantic control? We examined the overlap between brain regions involved in demanding syntactic processing and three networks: the SCN (generated above), the multiple-demand network (MDN) responsible for domain-general executive control, and regions supporting general semantic cognition that are not implicated in either form of control (semantic, not SCN/MDN). The MDN mask was generated by Fedorenko et al. (2013), and the general semantic cognition mask was derived from Jackson (2021).

Consistent with the demonstration above, demanding syntactic processing mostly fell within the SCN. However, there was additional recruitment of the left dorsal frontal cortices and left IPL, regions within the MDN. A smaller part of the pSTS/pMTG/pSTG fell within the semantic regions not involved in control (Figure 3). However, this area is within the SCN identified in Jackson (2021), it has simply failed to reach significance in the language-only version of this meta-analysis conducted here. Additionally, it is within the regions thought to cause semantic control impairments when damaged (Jefferies & Lambon Ralph, 2006;Thompson et al., 2018). Moreover, the key regions implicated in semantic cognition, yet not semantic control, such as the ATL and ventral angular gyrus, were not reliably activated for syntactic demands. Overall, 53.7% syntactic regions fell within the SCN, 19.4% within the MDN, and 12.7% within the semantic, not SCN/MDN regions (4345 voxels in total, Figure 3).

**Figure 3.**
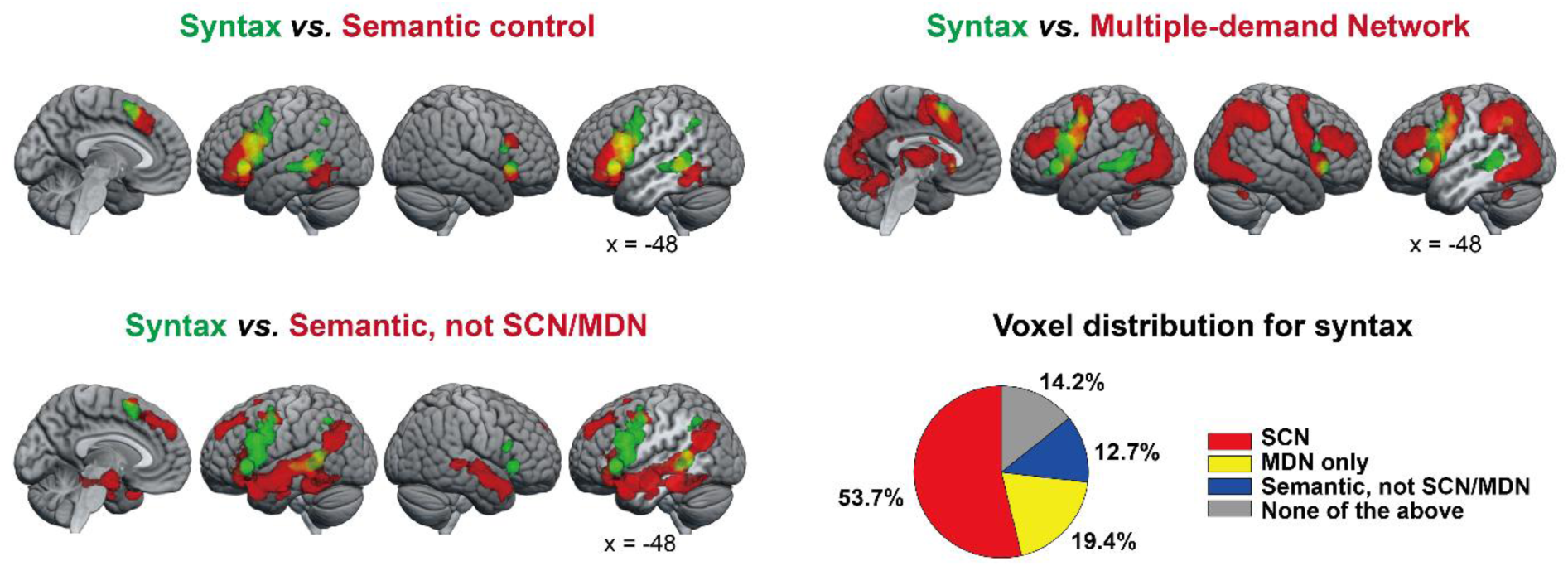
Top, left: overlap between syntax (green) and semantic control (blue). Overlap shown in cyan. Top, right: syntax (green) in the context of multiple-demand network (MDN) mask generated in Federenko et al., 2013 (blue). Overlap shown in cyan. Bottom, left: overlap between syntax (green) and general semantic cognition not implicated in the semantic control network (SCN) or MDN (semantic, not SCN/MDN, blue). The general semantic cognition mask was generated in Jackson (2021). Overlap shown in cyan. Bottom, right: voxel distribution of syntax within (i) the SCN. (ii) MDN not implicated in the SCN. (iii) semantic not SCN/MDN. Voxels outside these networks are also shown (none of the above). 4345 voxels in total.

### 3.4 Syntactic processing and semantic control regions in visual and auditory modalities

Next, we asked if the effect of input modality is the same for syntax and semantic control. More studies involved visual presentation for both syntactic and semantic control. Indeed, visual syntactic processing recruited a similar network to general syntactic processing, while auditory syntactic processing identified more circumscribed regions of left IFG (pars triangularis and pars opercularis), left pMTG/pSTS, and dmPFC (Figure 4 and Table 3), which were shared with the visual condition. Compared to auditory syntactic processing, visual syntactic processing showed greater activation in the left IPL and left IFG (pars triangularis and pars opercularis) extending to MFG (Figure 4 and Table 4).

**Figure 4.**
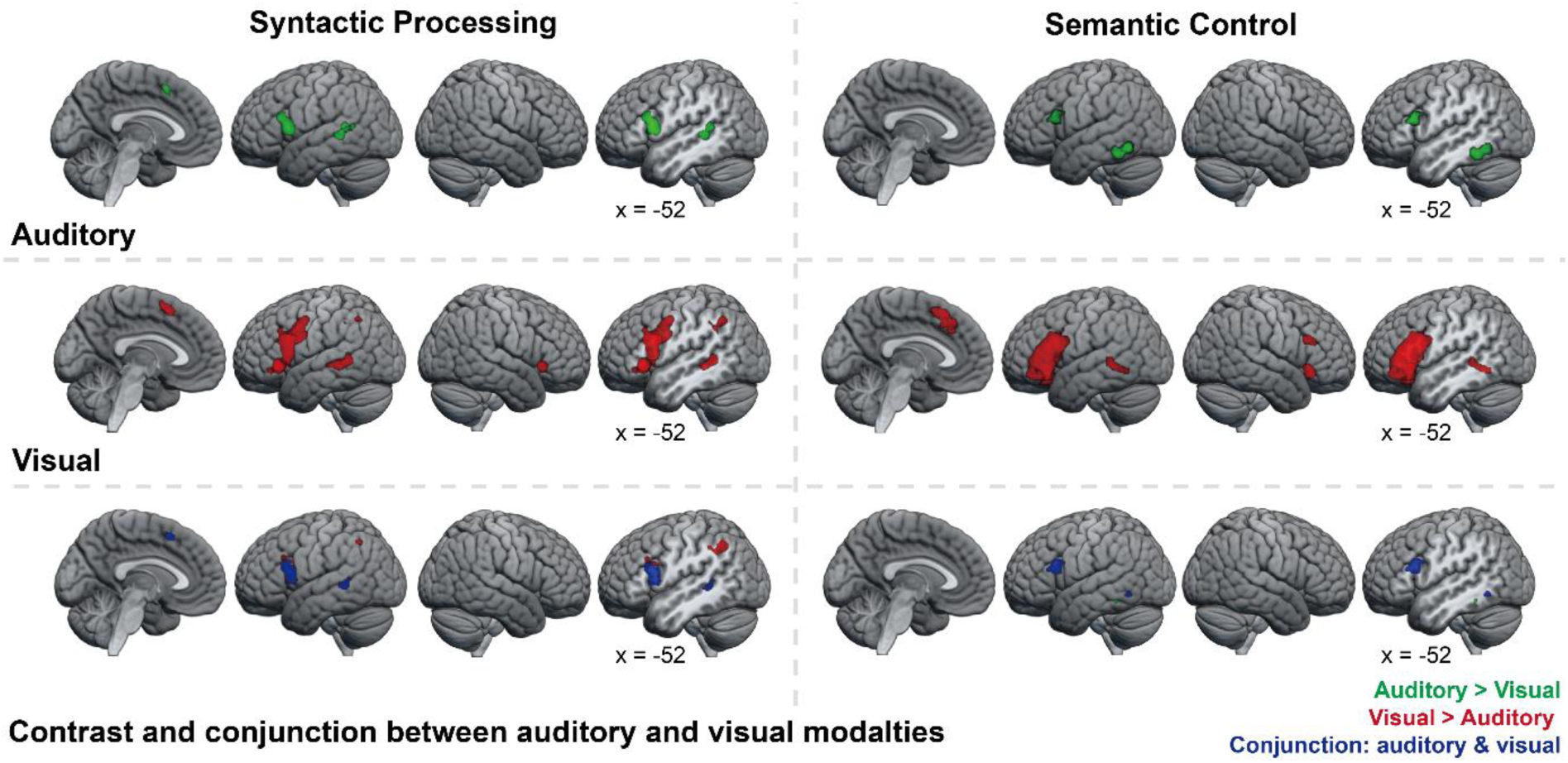
Top row: Activation likelihood estimation maps for auditory syntactic processing (left) and auditory semantic control (right). Second row: Activation likelihood estimation maps for visual syntactic processing (left) and visual semantic control (right). Bottom row: Contrast and conjunction analyses between auditory and visual modalities for syntactic processing (left) and semantic control (right), auditory > visual in green, visual > auditory in red, and conjunction of auditory and visual modalities in blue.

**Table 3.**
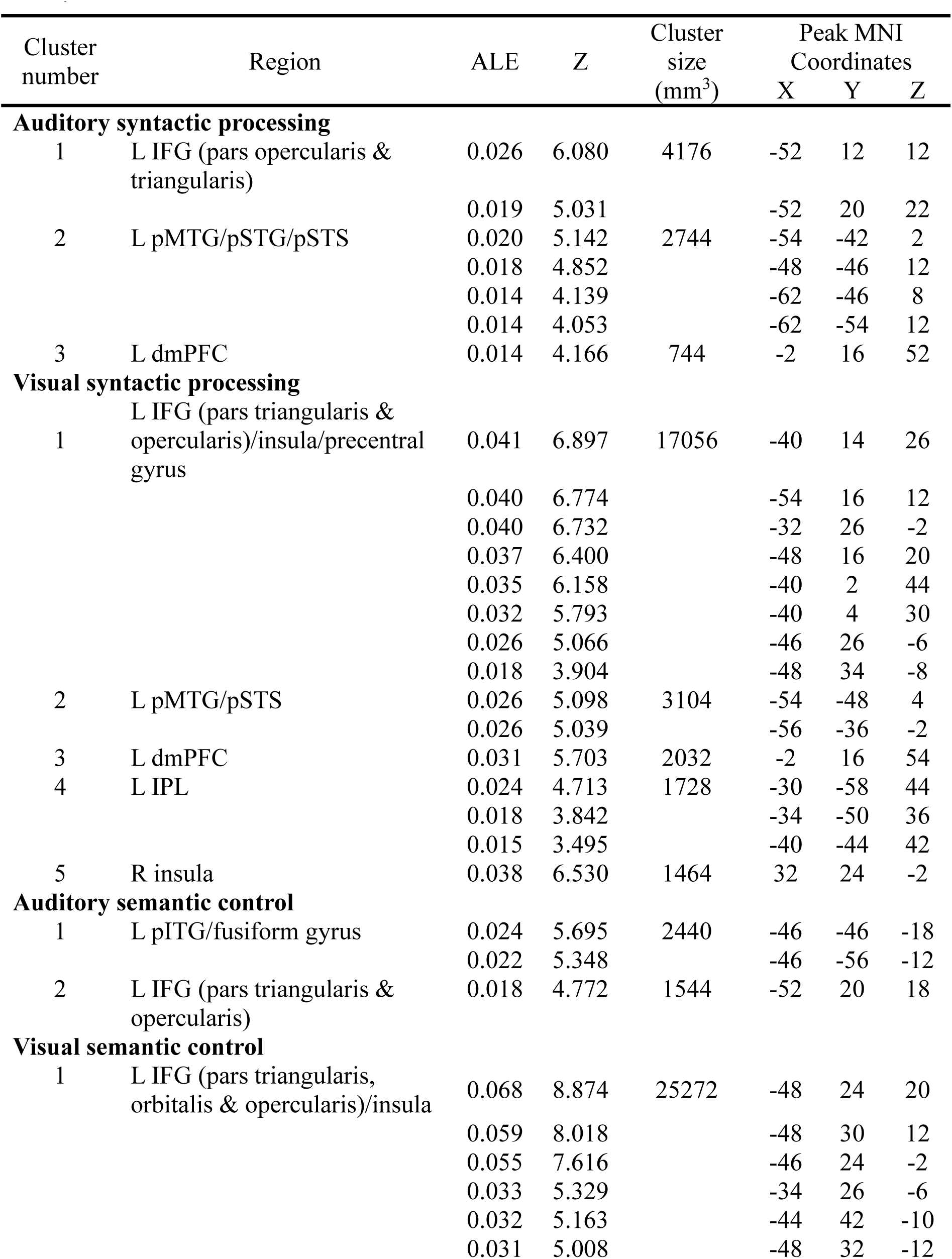
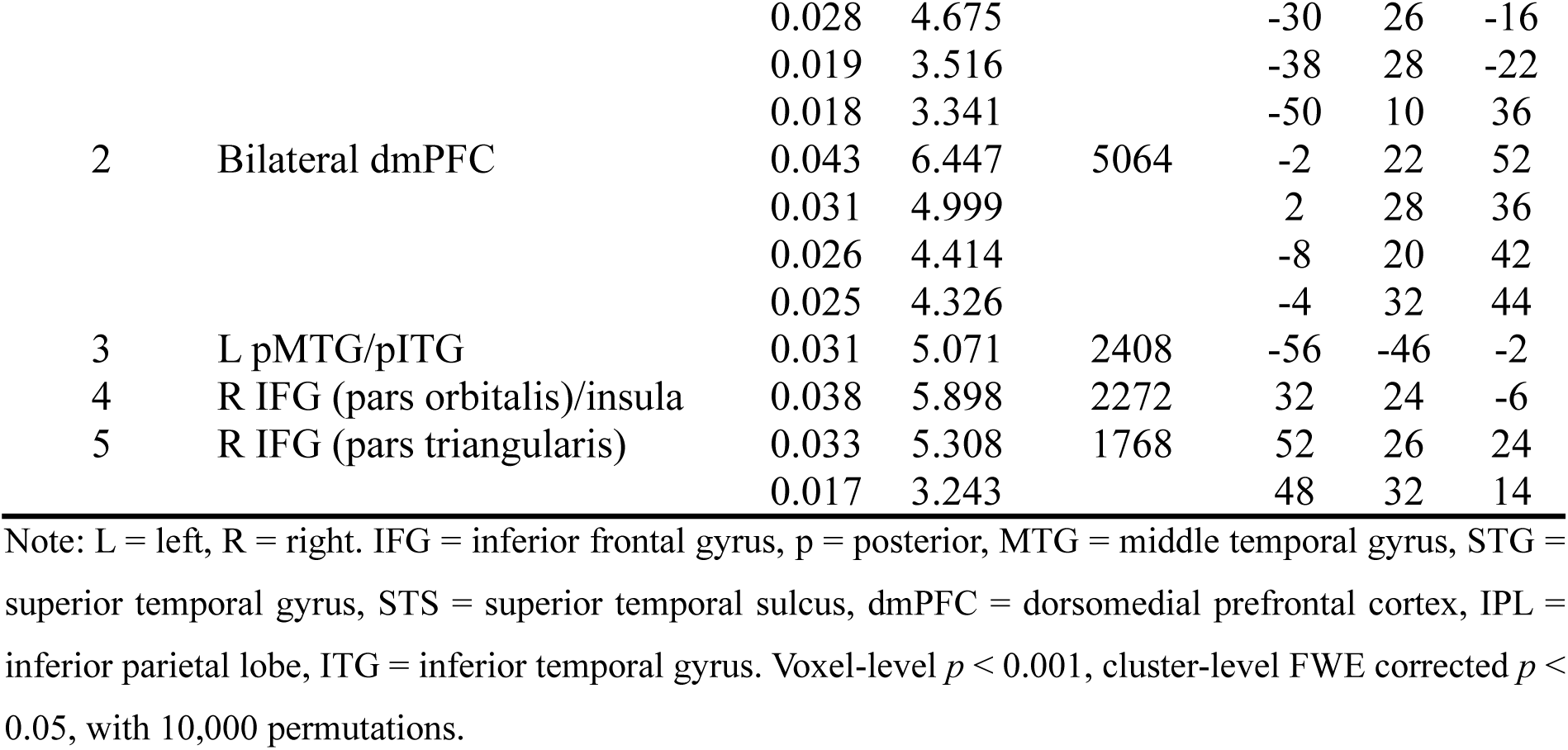
Activation likelihood estimation values for syntactic processing and semantic control in visual and auditory modalities.

**Table 4.** Contrast and conjunction analyses between auditory and visual modalities.

| Cluster number | Region | Cluster size (mm<sup>3</sup>) | Peak MNI Coordinates |  |  |
| --- | --- | --- | --- | --- | --- |
|  |  |  | X | Y | Z |
| **Syntactic processing** |  |  |  |  |  |
| *auditory syntactic processing > visual syntactic processing* |  |  |  |  |  |
| No significant results |  |  |  |  |  |
| *visual syntactic processing > auditory syntactic processing* |  |  |  |  |  |
| 1 | L IPL | 1328 | -33 | -56 | 45 |
|  |  |  | -37 | -52 | 36 |
| 2 | L IFG (pars triangularis & opercularis)/MFG | 288 | -42 | 24 | 32 |
|  |  |  | -37 | 15 | 26 |
|  |  |  | -36 | 10 | 24 |
| *Conjunction of auditory syntactic processing & visual syntactic processing* |  |  |  |  |  |
| 1 | L IFG (pars opercularis & triangularis) | 3032 | -52 | 12 | 12 |
|  |  |  | -52 | 20 | 22 |
| 2 | L pMTG/pSTS | 784 | -54 | -42 | 2 |
|  |  |  | -50 | -46 | 8 |
| 3 | L dmPFC | 496 | -2 | 16 | 52 |
| **Semantic control** |  |  |  |  |  |
| *Auditory semantic control > visual semantic control* |  |  |  |  |  |
| 1 | L pITG/fusiform gyrus | 56 | -43 | -44 | -15 |
| *Visual semantic control > auditory semantic control* |  |  |  |  |  |
| No significant results |  |  |  |  |  |
| *Conjunction of auditory semantic control & visual semantic control* |  |  |  |  |  |
| 1 | L IFG (pars triangularis & opercularis) | 1528 | -52 | 20 | 18 |
| 2 | L pITG | 120 | -48 | -58 | -10 |

Note: L = left, R = right. IPL = inferior parietal lobe, IFG = inferior frontal gyrus, MFG = middle frontal gyrus, p = posterior, MTG = middle temporal gyrus, STS = superior temporal sulcus, dmPFC = dorsomedial
prefrontal cortex, ITG = inferior temporal gyrus. Voxel-level $p < 0.001$ with 10,000 permutations, cluster volume $> 20 \text{ mm}^3$ .

We replicated the separation of semantic control conditions into auditory verbal and visual verbal stimuli from Jackson’s (2021) meta-analysis. As a greater number of studies were visual, more of the SCN was identified for visual than auditory semantic control (see Figure 4 and Table 3). The two modalities shared activation in the left prefrontal and posterior temporal cortex (IFG and pITG) (Figure 4 and Table 4), supporting the multimodal nature of these core semantic control regions. Despite this, significantly greater involvement was identified for a small region of pITG for auditory semantic control.

Both semantic control and syntactic processing were supported by modality-general regions of the inferior frontal and posterior temporal cortex. However, increasing syntactic demands for visually presented stimuli may also recruit modality-specific regions outside of (or on the edge of) semantic control areas, including inferior parietal cortex and posterior IFG. For both domains, the spatial extent of the visual network was larger than that of the auditory network, likely due to the smaller number of studies using auditory stimuli.

## 4 Discussion

Prior assessments of the brain regions engaged by semantic cognition and syntax fail to consider the crucial separation between semantic representation and control. This study used formal meta-analyses to identify whether demanding syntactic processing recruits the semantic control network. Remarkably, demanding syntactic processing engaged every region of the distributed SCN with the same pattern of lateralisation as semantic control, including IFG (in both hemispheres, yet particularly on the left), bilateral dmPFC, and left lateral posterior temporal cortex. Substantial overlap can be seen across the left IFG and posterior temporal cortex regions known to be necessary for semantic control (Jefferies & Lambon Ralph, 2006; Thompson et al., 2022). In contrast, key semantic representation areas were not recruited.

Despite this overlap, direct contrasts revealed some distinctions between the regions engaged by the two processes. There was a greater likelihood of activation for the anterior aspects of IFG and the posterior ITG for semantic control than demanding syntax. No regions outside of the SCN showed greater activation likelihood for demanding syntax than semantic control. However, a small cluster within the posterior IFG, bordering the precentral gyrus, did show this pattern. Additionally, demanding syntax consistently recruited a left IPL region, and the lateral frontal cluster extended more dorsally, although these regions were not identified as significantly more active in the contrast analysis.

Like controlled semantic processing, demanding syntactic processing engaged SCN regions irrespective of the modality of presentation (written versus spoken words). Yet unlike semantic control, demanding syntax also demonstrated modality-specific responses, with greater activation likelihood for visual than auditory stimuli in dorsal IFG/MFG and IPL. Thus, controlled processing of syntax appears to engage the multimodal SCN, perhaps as well as additional regions associated with domain-general executive control, particularly for visually presented stimuli.

The present study also considers the association between syntax and distinct components of semantic cognition: we separately visualise the overlap between demanding syntax and semantic control and the overlap between syntax and semantic regions not implicated in control. The overlap between syntax and semantics was strongly focused on the distributed SCN. The overlap identified within the pSTS/MTG may reflect control or language-related processes (Hickok & Poeppel, 2007; Hodgson et al., 2024; Jackson, 2021) but is not a key area for representing semantic content. Semantic representation areas, including the multimodal anterior temporal lobe (ATL) hub, were not implicated in demanding syntactic processing. This is consistent with the preservation of syntax in semantic dementia (Landin-Romero et al., 2016; Rochon et al., 2004). By separating semantic cognition into semantic control and representation components, we are able to explain both prior associations and dissociations with syntax in a systematic fashion (Hagoort & Indefrey, 2014; Rodd et al., 2015; Turker et al., 2023; Vigneau et al., 2006, 2011). Maintaining a separation between semantic representations and syntax allows orthogonal manipulation of meaning and sentence structure, for instance, enabling novel sentences to be easily generated when new vocabulary is learnt. Yet, the SCN acts as a shared resource to support both demanding syntactic and semantic processing.

Why does demanding syntax engage the SCN? One possibility is methodological: some apparent syntactic effects may reflect incidental engagement of semantic control processes, since sentences are occasionally used as stimuli in semantic control studies. Yet this explanation seems insufficient, as most semantic control tasks use isolated words or pictures lacking syntactic structure, and the same regions remain critical (Grindrod et al., 2008; Krieger-Redwood et al., 2015; Snyder et al., 2011; Tobia & Madan, 2017). A more compelling account is that demanding syntactic and semantic tasks share a common computational goal – namely, deriving context-sensitive meaning. The function of syntax itself may be to constrain and clarify possible interpretations, such as distinguishing whether *the cat chased the dog* or *the dog chased the cat*. When syntactic complexity increases, semantic prediction becomes less reliable, requiring active resolution among competing interpretations. This selective retrieval and inhibition of context-appropriate versus inappropriate meanings is the hallmark of semantic control (Chiou et al., 2018; Jefferies, 2013; Lambon Ralph et al., 2017). Consequently, both syntactic and semantic challenges may engage overlapping SCN regions because each demands the extraction and integration of meaning that is flexible yet contextually constrained.

While demanding syntactic processing engaged the SCN, consistent activity was also found in IPL and more dorsal frontal cortices. This aligns with a previous meta-analysis showing higher activation likelihood for demanding syntax than demanding semantics (i.e., semantic control) in the left IPL (Rodd et al., 2015). Lesion mapping work has also shown that syntactic comprehension deficits in aphasia are more strongly associated with damage to left posterior temporal-parietal cortex (Matchin et al., 2023, 2024). Additionally, a posterior IFG region bordering the precentral gyrus showed greater involvement for syntax. These regions are not uniquely associated with syntax, but form part of another network, known as the multiple-demand network (MDN), which underpins executive control across diverse cognitive domains, stimulus modalities and task processes (Camilleri et al., 2018; Duncan, 2010). The MDN comprises bilateral frontoparietal regions, including dmPFC, premotor cortex, MFG, insula, and IPL (Assem et al., 2020; Duncan, 2010; Fedorenko et al., 2013). While the parietal cortex is not reliably activated across studies of semantic control (Jackson, 2021), inferior parietal cortex is a core region of the MDN (Assem et al., 2020; Fedorenko et al., 2013) and the domain-general precentral gyrus and dorsolateral prefrontal cortex surround the semantic IFG. MDN engagement for demanding syntax would not be surprising, given the network’s role in demanding processing across domains. This converging evidence suggests that, in addition to the SCN, demanding syntax also engages some regions of the MDN.

The presence of meaningful stimuli typically down-regulates activity in the MDN while increasing engagement of the SCN (Hodgson et al., 2024). These networks support distinct forms of control: the SCN facilitates context-sensitive retrieval and selection of meaning, whereas the MDN supports domain-general executive demands such as maintaining and manipulating information in working memory (Gao et al., 2021). The present results suggest that demanding syntactic processing draws on both forms of control. Complex syntactic structures not only require context-sensitive interpretation, invoking the SCN, but also place heavier demands on working memory, engaging elements of the MDN. Consistent with this, tasks emphasising phonological rather than semantic control preferentially recruit the MDN (Hodgson et al., 2021; Noonan et al., 2013). Demanding syntax may therefore co-activate SCN regions needed to manage meaning and MDN systems that support structural integration of the elements of a sentence. Importantly, we found no evidence for additional, syntax-specific regions beyond these established control networks.

The SCN showed equivalent responses to visual and auditory stimuli whether performing demanding syntactic or semantic processing. Indeed, semantic aphasia patients have deficits across both visual and auditory semantic control (Thompson & Jefferies, 2013). In addition, demanding syntax showed modality-specific responses in MDN regions, including left dorsal IFG/MFG and IPL. These regions showed a greater activation likelihood for visual than auditory stimuli. One possible explanation is a visual preference within the MDN. Previous studies have demonstrated sensory biases within the MDN, including a visually-biased region in the left inferior precentral sulcus bordering IFG and MFG (Assem et al., 2022; Mayer et al., 2017; Noyce et al., 2017; Tobyne et al., 2017). An alternative explanation is that these differences could arise from participants re-reading the visual stimuli to aid comprehension, while auditory stimuli are fleeting and cannot be revisited.

In conclusion, the present study used formal meta-analysis to demonstrate that syntactic processing relies on the semantic control network, encompassing left-dominant frontotemporal regions including IFG, insula, left posterior temporal cortex, and bilateral dmPFC. Beyond the SCN, demanding syntactic processing may recruit additional regions within the multiple-demand network (MDN), including the left posterior IFG and left IPL, yet is distinct from core semantic representation areas. Furthermore, for demanding syntax, SCN regions showed multimodal responses to both visual and auditory stimuli, whereas MDN regions (i.e., left IFG/MFG and left IPL) showed a greater activation likelihood for visual than auditory stimuli. Dissociating semantic control and representation is crucial to understand the relationship between semantic cognition and syntax.

## Data and code availability

All data are provided in the Supplementary Materials, and the syntax meta-analysis results are available online at https://github.com/QianwenChang/Syntax_SemanticControl_Meta#.

