## Supplementary Table 1 for "Syntactic Processing Engages the Semantic Control Network"

**Supplementary Table 1**. The 66 experiments included in the meta-analysis of syntactic processing

| Study | DOI | N | Modality | Syntactic  manipulation | Space | Contrast | X | Y | Z |
| --- | --- | --- | --- | --- | --- | --- | --- | --- | --- |
| Bahlmann et al. (2007) | 10.1002/hbm.20318 | 12 | visual | complexity | MNI | non-canonical > canonical word order (subject first > object first) | -40 | -52 | 28 |
|  |  |  |  |  |  |  | -44 | 24 | 28 |
| Ben-Shachar et al. (2003) | 10.1111/1467-9280.01459 | 11 | auditory | complexity | TAL | transformational > non-transformational sentence | -47 | 18 | 7 |
|  |  |  |  |  |  |  | -37 | -40 | 20 |
| Bornkessel-Schlesewsky et al. (2009) | 10.1016/j.bandl.2009.09.004 | 28 | visual | complexity | TAL | object-subject > subject-object sentence | -53 | 11 | 5 |
|  |  |  |  |  |  |  | -34 | 23 | 5 |
|  |  |  |  |  |  |  | -40 | 5 | 32 |
|  |  |  |  |  |  |  | 31 | 20 | 6 |
|  |  |  |  |  |  |  | -7 | 23 | 41 |
|  |  |  |  |  |  |  | -32 | -58 | 41 |
|  |  |  |  |  |  |  | -14 | 1 | 18 |
| Caplan et al. (1998) | 10.1006/nimg.1998.0412 | 16 | auditory | complexity | TAL | cleft object > cleft subject sentence | -52 | 18 | 24 |
| Caplan 1998 | 10.1162/089892998562843 | 8 | visual | complexity | TAL | center-embedded > right-branching sentence | 10 | 6 | 52 |
|  |  |  |  |  |  |  | -2 | 6 | 40 |
|  |  |  |  |  |  |  | -42 | 18 | 24 |
| Caplan 2000 | 10.1002/(SICI)1097-0193(200002)9:2<65::AID-HBM1>3.0.CO;2-4 | 11 | visual | complexity | TAL | Center-embedded > right-branching clauses | -46 | 36 | 4 |
|  |  |  |  |  |  |  | -14 | -20 | 4 |
|  |  |  |  |  |  |  | -10 | -36 | 40 |
|  |  |  |  |  |  |  | 0 | 56 | 8 |
| Caplan et al.. (2008) | 10.1162/jocn.2008.20044 | 16 | visual | complexity | MNI | object-extracted relative clause > subject-extracted relative clause (plausible unconstrained sentences) | -52 | -44 | -2 |
|  |  |  |  |  |  |  | -46 | -56 | 16 |
|  |  |  |  |  |  |  | -62 | -18 | -1 |
|  |  |  |  |  |  |  | -38 | 30 | 0 |
|  |  |  |  |  |  |  | -44 | 34 | -14 |
|  |  |  |  |  |  |  | -46 | 24 | -8 |
|  |  |  |  |  |  |  | -54 | 18 | 28 |
|  |  |  |  |  |  |  | -46 | 14 | 18 |
|  |  |  |  |  |  | object-extracted relative clause > subject-extracted relative clause (plausible constrained sentences) | -24 | -76 | 32 |
|  |  |  |  |  |  |  | -28 | -84 | 22 |
|  |  |  |  |  |  |  | -24 | -86 | 32 |
|  |  |  |  |  |  |  | -58 | -46 | 2 |
|  |  |  |  |  |  |  | -50 | -60 | 12 |
|  |  |  |  |  |  |  | -50 | -42 | -4 |
| Carreiras 2015 | 10.1016/j.neuroimage.2015.06.075 | 32 | visual | violation | MNI | number disagreement > agreement (subject–verb and determiner–noun) | 32 | 26 | -2 |
|  |  |  |  |  |  |  | 28 | 20 | 4 |
|  |  |  |  |  |  |  | -36 | -42 | 42 |
|  |  |  |  |  |  |  | -28 | 24 | 0 |
|  |  |  |  |  |  |  | -50 | 10 | 26 |
|  |  |  |  |  |  |  | -42 | 20 | -4 |
|  |  |  |  |  |  |  | -40 | 14 | 28 |
|  |  |  |  |  |  |  | -46 | -2 | 48 |
|  |  |  |  |  |  |  | -6 | 4 | 62 |
| Chen 2006 | 10.1016/S0010-9452(08)70397-6 | 12 | visual | complexity | TAL | Object relative > subject relative clause | -60 | 16 | 14 |
|  |  |  |  |  |  |  | -46 | -68 | 30 |
|  |  |  |  |  |  |  | -41 | -43 | 40 |
|  |  |  |  |  |  |  | -53 | 9 | 40 |
|  |  |  |  |  |  |  | 36 | -61 | 38 |
|  |  |  |  |  |  |  | 25 | -56 | 48 |
| Constable et al. (2004) | [10.1016/j.neuroimage.2004.01.001](https://dx.doi.org/10.1016/j.neuroimage.2004.01.001) | 20 | visual & auditory | complexity | TAL | object relative > subject relative sentence | -51 | -58 | 3 |
|  |  |  |  |  |  |  | -49 | 11 | 13 |
|  |  |  |  |  |  |  | -36 | -64 | 31 |
|  |  |  |  |  |  |  | -36 | 4 | 46 |
|  |  |  |  |  |  |  | -2 | 6 | 33 |
|  |  |  |  |  |  |  | -3 | -24 | 15 |
|  |  |  |  |  |  |  | 44 | 6 | 2 |
|  |  |  |  |  |  |  | 43 | 14 | 23 |
| Cooke et al. (2001) | 10.1002/hbm.10006 | 7 | visual | complexity | TAL | object- > subject-relative center-embedded clause (short antecedent-gap distance) | -48 | -68 | -8 |
|  |  |  |  |  |  |  | -4 | -92 | -8 |
|  |  |  |  |  |  |  | 28 | -68 | -20 |
|  |  |  |  |  |  | object- > subject-relative center-embedded clause (long antecedent-gap distance) | -40 | -76 | -4 |
|  |  |  |  |  |  |  | -32 | -20 | -20 |
|  |  |  |  |  |  |  | 36 | -40 | -12 |
|  |  |  |  |  |  |  | 16 | -92 | -12 |
| Cooke et al. (2006) | 10.1016/j.bandl.2005.07.072 | 15 | visual | violation | TAL | Inflectional morphology violation > correct sentence (early phase) | -60 | 8 | 24 |
|  |  |  |  |  |  |  | -60 | -52 | 8 |
|  |  |  |  |  |  |  | 4 | 16 | 48 |
|  |  |  |  |  |  | Grammatical category violation > correct sentence (early phase) | -44 | 12 | 24 |
|  |  |  |  |  |  |  | -60 | -52 | 4 |
|  |  |  |  |  |  |  | -12 | 8 | 52 |
|  |  |  |  |  |  |  | 48 | 8 | 32 |
|  |  |  |  |  |  |  | 36 | -64 | 44 |
|  |  |  |  |  |  | Grammatical category violation > correct sentence (late phase) | -48 | 12 | 0 |
|  |  |  |  |  |  |  | -44 | -52 | 0 |
|  |  |  |  |  |  | Transitivity violation > correct sentence (early phase) | -52 | 0 | 40 |
|  |  |  |  |  |  |  | 4 | 8 | 52 |
|  |  |  |  |  |  |  | 48 | 8 | 40 |
|  |  |  |  |  |  | Transitivity violation > correct sentence (late phase) | -48 | 8 | 12 |
| Feng et al. (2015) | 10.1016/j.jneuroling.2014.09.002 | 18 | visual | complexity | MNI | Passive > active sentences | -48 | 26 | 0 |
|  |  |  |  |  |  |  | -56 | -44 | 12 |
|  |  |  |  |  |  |  | -44 | -2 | 46 |
|  |  |  |  |  |  |  | -62 | 0 | -14 |
|  |  |  |  |  |  |  | -18 | -48 | 2 |
|  |  |  |  |  |  |  | -34 | -22 | 24 |
|  |  |  |  |  |  |  | 32 | 0 | 20 |
|  |  |  |  |  |  |  | 24 | -28 | -26 |
| Fiebach et al. (2005) | 10.1002/hbm.20070 | 14 | visual | complexity | TAL | object questions with long > short dislocation | -44 | 21 | 11 |
|  |  |  |  |  |  |  | -45 | 21 | 10 |
|  |  |  |  |  |  |  | -46 | 17 | 4 |
|  |  |  |  |  |  |  | -54 | 7 | 28 |
|  |  |  |  |  |  |  | -54 | -27 | -1 |
|  |  |  |  |  |  |  | -52 | -46 | 6 |
|  |  |  |  |  |  |  | 45 | -18 | -3 |
|  |  |  |  |  |  |  | -18 | -18 | 12 |
| Friederici et al. (2006) | [10.1093/cercor/bhj106](https://dx.doi.org/10.1093/cercor/bhj106) | 13 | visual | violation | TAL | grammatically incorrect > correct sentence | -43 | -23 | 52 |
|  |  |  |  |  |  |  | 46 | -43 | 52 |
|  |  |  |  |  |  |  | 34 | -65 | 44 |
|  |  |  |  |  |  |  | -31 | -70 | -17 |
| Friederici et al. (2010) | [10.1002/hbm.20878](https://dx.doi.org/10.1002/hbm.20878) | 17 | auditory | violation | MNI | grammatically incorrect > correct sentence | -54 | -16 | 4 |
|  |  |  |  |  |  |  | -60 | -42 | 6 |
|  |  |  |  |  |  |  | -62 | 18 | 16 |
|  |  |  |  |  |  |  | -12 | -28 | 2 |
|  |  |  |  |  |  |  | 44 | -20 | 6 |
| Grewe et al. (2005) | 10.1002/hbm.20154 | 16 | visual | complexity | TAL | permuted non-pronominal (N-OS) > non-permuted non-pronominal (N-SO) | -32 | 20 | 3 |
|  |  |  |  |  |  |  | -52 | 14 | 15 |
|  |  |  |  |  |  |  | -2 | 32 | 30 |
|  |  |  |  |  |  |  | -38 | 8 | 38 |
|  |  |  |  |  |  |  | 44 | 26 | 18 |
|  |  |  |  |  |  |  | 38 | 20 | 6 |
|  |  |  |  |  |  |  | 46 | 11 | 9 |
| Grewe 2007 | 10.1016/j.neuroimage.2006.11.045 | 19 | visual | complexity | TAL | object > subject-initial sentences | -53 | 10 | 15 |
|  |  |  |  |  |  |  | -20 | -86 | 24 |
|  |  |  |  |  |  |  | 28 | -8 | 15 |
| Herrmann et al. (2012) | 10.1002/hbm.21235 | 25 | auditory | violation | MNI | syntactically incorrect > correct | -54 | 8 | 10 |
|  |  |  |  |  |  |  | -60 | -22 | -2 |
|  |  |  |  |  |  |  | -54 | 5 | -14 |
|  |  |  |  |  |  |  | 57 | -28 | 1 |
|  |  |  |  |  |  |  | 60 | -4 | -8 |
|  |  |  |  |  |  |  | -45 | -22 | 1 |
|  |  |  |  |  |  |  | 48 | -22 | 7 |
| Husband et al. (2011) | 10.1162/jocn_a_00040 | 19 | visual | violation | TAL | ungrammatical > grammatical sentence | -45 | -65 | 30 |
|  |  |  |  |  |  |  | -46 | 1 | -27 |
|  |  |  |  |  |  |  | -29 | -39 | 57 |
|  |  |  |  |  |  |  | -6 | -15 | 50 |
|  |  |  |  |  |  |  | -45 | -65 | 29 |
|  |  |  |  |  |  |  | -6 | 58 | 23 |
|  |  |  |  |  |  |  | 45 | -65 | 32 |
|  |  |  |  |  |  |  | 45 | -65 | 28 |
| Iwabuchi 2020 | 10.1016/j.jneuroling.2020.100893 | 23 | visual | complexity | MNI | scrambled subject-second > canonical subject-first sentence | -3 | 17 | 55 |
|  |  |  |  |  |  |  | 33 | 23 | -8 |
|  |  |  |  |  |  |  | -42 | 14 | 25 |
|  |  |  |  |  |  |  | -30 | 26 | -2 |
|  |  |  |  |  |  |  | -39 | 2 | 43 |
|  |  |  |  |  |  |  | -30 | -61 | 43 |
| Kambara et al. (2013) | 10.1016/j.langsci.2012.07.003 | 38 | visual | violation | TAL | syntactically violation > correct sentence | -59 | -33 | 38 |
|  |  |  |  |  |  |  | 42 | -52 | 58 |
|  |  |  |  |  |  |  | 6 | -65 | 45 |
|  |  |  |  |  |  |  | 24 | -33 | -28 |
| Koizumi et al. (2016) | 10.3389/fpsyg.2016.01541 | 16 | visual | complexity | MNI | complex Subject-Verb-Object > simple Verb-Object-Subject | -42 | 44 | 1 |
| Kristensen et al. (2013) | 10.1016/j.jneuroling.2012.05.001 | 21 | visual | complexity | MNI | object-initial > subject-initial sentence | -6 | 10 | 58 |
|  |  |  |  |  |  |  | -54 | 14 | 10 |
|  |  |  |  |  |  |  | -38 | -2 | 58 |
|  |  |  |  |  |  |  | 10 | 8 | 2 |
|  |  |  |  |  |  |  | 32 | 22 | -2 |
|  |  |  |  |  |  |  | -52 | -50 | 4 |
|  |  |  |  |  |  |  | -28 | -60 | 46 |
|  |  |  |  |  |  |  | -14 | -68 | 56 |
|  |  |  |  |  |  |  | -14 | -10 | 6 |
|  |  |  |  |  |  |  | -10 | 10 | 4 |
| Kristensen et al. (2014) | 10.1162/jocn_a_00681 | 32 | auditory | complexity | MNI | object-initial > subject-initial sentence in the main task | 0 | 18 | 44 |
|  |  |  |  |  |  |  | -14 | -74 | -28 |
|  |  |  |  |  |  |  | 32 | 28 | -6 |
| Kroczek et al. (2020) | 10.1093/texcom/tgaa021 | 28 | auditory | complexity | MNI | complex OSV > easy SOV sentence | -54 | 11 | 5 |
|  |  |  |  |  |  |  | -42 | 2 | 53 |
|  |  |  |  |  |  |  | -54 | -40 | 2 |
| Kunert et al. (2015) | 10.1371/journal.pone.0141069 | 19 | auditory | complexity | MNI | object-extracted relative clause > subject-extracted relative clause | -54 | 18 | 28 |
| Kuperberg et al. (2000) | 10.1162/089892900562138 | 9 | auditory | violation | TAL | syntactically violated > normal sentence | -26 | -14 | -13 |
|  |  |  |  |  |  |  | 12 | -31 | -2 |
|  |  |  |  |  |  |  | -17 | -33 | -2 |
|  |  |  |  |  |  |  | -38 | -42 | -2 |
|  |  |  |  |  |  |  | -35 | -69 | 9 |
|  |  |  |  |  |  |  | -49 | -53 | 9 |
|  |  |  |  |  |  |  | 49 | -14 | -2 |
|  |  |  |  |  |  |  | -46 | -47 | 15 |
|  |  |  |  |  |  |  | 46 | 22 | -2 |
|  |  |  |  |  |  |  | 20 | -53 | 4 |
|  |  |  |  |  |  |  | 20 | -53 | -2 |
|  |  |  |  |  |  |  | -3 | -78 | -2 |
|  |  |  |  |  |  |  | -9 | -53 | 4 |
|  |  |  |  |  |  |  | -14 | -50 | 9 |
|  |  |  |  |  |  |  | 14 | -50 | 15 |
|  |  |  |  |  |  |  | 9 | -56 | -7 |
|  |  |  |  |  |  |  | -12 | -22 | 9 |
| Kuperberg 2008 | 10.1016/j.neuroimage.2007.10.009 | 16 | visual | violation | TAL | syntactic violated > normal sentence | -49 | 3 | 7 |
|  |  |  |  |  |  |  | 29 | 19 | 5 |
|  |  |  |  |  |  |  | -45 | -9 | 41 |
|  |  |  |  |  |  |  | 52 | 3 | 33 |
|  |  |  |  |  |  |  | -19 | 33 | 29 |
|  |  |  |  |  |  |  | 20 | 39 | 21 |
|  |  |  |  |  |  |  | -56 | -31 | 38 |
|  |  |  |  |  |  |  | -51 | -19 | 46 |
|  |  |  |  |  |  |  | -39 | -4 | 14 |
|  |  |  |  |  |  |  | 36 | -3 | 14 |
|  |  |  |  |  |  |  | -53 | -20 | -1 |
|  |  |  |  |  |  |  | 58 | -19 | 3 |
|  |  |  |  |  |  |  | -6 | 37 | -5 |
|  |  |  |  |  |  |  | 6 | 49 | -8 |
|  |  |  |  |  |  |  | -11 | -6 | 47 |
|  |  |  |  |  |  |  | 9 | -2 | 52 |
|  |  |  |  |  |  |  | -35 | -42 | -6 |
|  |  |  |  |  |  |  | 35 | -38 | -9 |
|  |  |  |  |  |  |  | -32 | -92 | 17 |
|  |  |  |  |  |  |  | 25 | -80 | 36 |
|  |  |  |  |  |  |  | -12 | 11 | -4 |
|  |  |  |  |  |  |  | 28 | -15 | 8 |
| Lee and Newman (2010) | 10.1002/hbm.20845 | 18 | visual | complexity | MNI | Object-relative > co-joined active sentence in the sentence reading phase | -42 | 4 | 28 |
|  |  |  |  |  |  | Object-relative > co-joined active sentence in the probe phase | -46 | 30 | -6 |
|  |  |  |  |  |  |  | 46 | 15 | 34 |
|  |  |  |  |  |  |  | -38 | 2 | 60 |
|  |  |  |  |  |  |  | -32 | 20 | -14 |
|  |  |  |  |  |  |  | -2 | 8 | 60 |
|  |  |  |  |  |  |  | -62 | -34 | 2 |
|  |  |  |  |  |  |  | -28 | -56 | 40 |
|  |  |  |  |  |  |  | 38 | -58 | 38 |
|  |  |  |  |  |  |  | 0 | -70 | 42 |
|  |  |  |  |  |  |  | 8 | -74 | -30 |
|  |  |  |  |  |  |  | 40 | -64 | -36 |
| Lee et al. (2016) | [10.1016/j.heares.2015.12.008](https://dx.doi.org/10.1016/j.heares.2015.12.008) | 26 | auditory | complexity | MNI | object-relative > subject relative | -60 | -55 | 14 |
|  |  |  |  |  |  |  | -12 | -13 | 8 |
|  |  |  |  |  |  |  | -48 | 17 | 20 |
|  |  |  |  |  |  |  | 33 | 29 | 2 |
|  |  |  |  |  |  |  | 18 | -73 | -28 |
|  |  |  |  |  |  |  | -39 | -70 | -34 |
|  |  |  |  |  |  |  | -9 | -79 | -28 |
|  |  |  |  |  |  |  | -3 | 17 | 53 |
|  |  |  |  |  |  |  | -6 | 38 | 44 |
|  |  |  |  |  |  |  | -9 | 20 | 32 |
|  |  |  |  |  |  |  | 39 | -1 | 50 |
|  |  |  |  |  |  |  | 48 | 23 | 23 |
|  |  |  |  |  |  |  | 45 | 5 | 38 |
|  |  |  |  |  |  |  | -60 | -7 | 14 |
|  |  |  |  |  |  |  | -48 | -13 | 20 |
|  |  |  |  |  |  |  | 24 | 65 | 14 |
|  |  |  |  |  |  |  | 42 | 56 | 2 |
|  |  |  |  |  |  |  | 36 | 59 | 14 |
|  |  |  |  |  |  |  | 63 | -7 | 17 |
|  |  |  |  |  |  |  | 66 | -19 | 26 |
|  |  |  |  |  |  |  | 63 | -4 | 29 |
| Lee 2018 | 10.1523/eneuro.0263-17.2018 | 35 | auditory | complexity | MNI | Object-relative > subject-relative center-embedded clause | -48 | -46 | 11 |
|  |  |  |  |  |  |  | -63 | -46 | 8 |
|  |  |  |  |  |  |  | -54 | -43 | 2 |
| Lee 2023 | 10.1016/j.jneuroling.2023.101126 | 31 | visual | complexity | MNI | topicalization > baseline sentence | -42 | 5 | 26 |
|  |  |  |  |  |  | object relative clause > baseline sentence | -51 | 29 | 2 |
|  |  |  |  |  |  |  | -57 | -28 | -2 |
|  |  |  |  |  |  |  | 66 | -13 | -6 |
|  |  |  |  |  |  |  | 63 | -1 | 26 |
|  |  |  |  |  |  | subject-relative clause > baseline sentence | -45 | -37 | -2 |
|  |  |  |  |  |  |  | -39 | 2 | 42 |
|  |  |  |  |  |  |  | 54 | -28 | -2 |
|  |  |  |  |  |  |  | -39 | -52 | -22 |
| Makuuchi et al. (2013) | 10.1093/cercor/bhs058 | 22 | visual | complexity | MNI | syntactic movement distance | -36 | 6 | 33 |
|  |  |  |  |  |  |  | -51 | 15 | 18 |
|  |  |  |  |  |  |  | -33 | -51 | 36 |
|  |  |  |  |  |  |  | 9 | -66 | 42 |
|  |  |  |  |  |  |  | -54 | -36 | -6 |
|  |  |  |  |  |  |  | -21 | -15 | 6 |
|  |  |  |  |  |  |  | 33 | -45 | 39 |
|  |  |  |  |  |  |  | 45 | 21 | 21 |
| Matchin et al. (2014) | 10.1016/j.bandl.2014.09.001 | 26 | auditory | complexity | TAL | syntactically long-distance dependency > short-distance dependency | -50 | 16 | 25 |
| Matchin 2016 | 10.3389/fpsyg.2016.00241 | 20 | visual | complexity | TAL | Passive > active sentences | -32 | -36 | 51 |
| Meltzer et al. (2010) | 10.1093/cercor/bhp249 | 24 | auditory | complexity | TAL | object-embedded clause > subject-embedded clause | -45 | 12 | 13 |
| Meyer 2012 | 10.1016/j.neuroimage.2012.05.052 | 24 | auditory | complexity | MNI | object-first > subject-first sentence | -54 | 14 | 13 |
| Nakagawa 2022 | [10.3389/fnhum.2021.753245](https://dx.doi.org/10.3389/fnhum.2021.753245) | 30 | visual | complexity | MNI | Double Object > Prepositional Object  structures | -46 | 50 | 8 |
|  |  |  |  |  |  |  | -48 | 42 | 16 |
|  |  |  |  |  |  |  | 0 | 16 | 54 |
|  |  |  |  |  |  |  | 4 | 22 | 46 |
|  |  |  |  |  |  |  | 56 | 16 | 26 |
|  |  |  |  |  |  |  | -48 | -36 | 42 |
|  |  |  |  |  |  |  | -44 | -44 | 54 |
|  |  |  |  |  |  |  | -48 | 26 | 36 |
|  |  |  |  |  |  |  | -46 | 6 | 22 |
|  |  |  |  |  |  |  | -54 | 20 | 26 |
|  |  |  |  |  |  |  | -40 | 4 | 30 |
|  |  |  |  |  |  |  | -58 | 16 | 16 |
| Newman et al. (2010) | 10.1016/j.bandl.2010.02.001 | 20 | visual | complexity | MNI | object-relative > co-joined sentence | -40 | 14 | 24 |
|  |  |  |  |  |  |  | -30 | 22 | 0 |
|  |  |  |  |  |  |  | -58 | -36 | 2 |
| Nieuwland et al. (2012) | 10.1002/hbm.21377 | 20 | visual | violation | MNI | case violation > correct | 0 | -30 | 28 |
|  |  |  |  |  |  |  | -8 | 0 | 36 |
|  |  |  |  |  |  |  | -4 | -28 | 46 |
|  |  |  |  |  |  |  | -18 | -68 | 32 |
|  |  |  |  |  |  |  | 4 | -70 | 40 |
|  |  |  |  |  |  |  | -4 | -70 | 44 |
|  |  |  |  |  |  |  | 50 | -44 | 40 |
|  |  |  |  |  |  |  | 60 | -38 | 32 |
|  |  |  |  |  |  |  | 46 | -58 | 46 |
|  |  |  |  |  |  |  | -42 | -46 | 42 |
|  |  |  |  |  |  |  | -52 | -46 | 44 |
|  |  |  |  |  |  |  | -60 | -34 | 36 |
|  |  |  |  |  |  | number agreement violation > correct | 36 | -52 | 42 |
|  |  |  |  |  |  |  | 42 | -44 | 38 |
|  |  |  |  |  |  |  | -44 | 32 | 34 |
|  |  |  |  |  |  |  | -44 | 54 | 8 |
|  |  |  |  |  |  |  | -44 | 48 | 24 |
|  |  |  |  |  |  |  | -42 | -46 | 42 |
|  |  |  |  |  |  |  | -46 | -36 | 36 |
|  |  |  |  |  |  |  | 44 | 46 | 16 |
|  |  |  |  |  |  |  | 50 | 40 | 20 |
|  |  |  |  |  |  |  | 42 | 50 | 6 |
| Obleser et al. (2011) | 10.1016/j.neuroimage.2011.03.035 | 16 | auditory | complexity | MNI | correlation with increasingly complex syntactic structure | -50 | 16 | -20 |
|  |  |  |  |  |  |  | -64 | -54 | 10 |
|  |  |  |  |  |  |  | -52 | 12 | 14 |
|  |  |  |  |  |  |  | 44 | 12 | 18 |
|  |  |  |  |  |  |  | 52 | 16 | 2 |
| Obleser et al. (2011) | 10.1016/j.neuroimage.2011.03.035 | 14 | auditory | complexity | MNI | correlation with increasingly complex syntactic structure | -48 | 10 | 18 |
| Ogawa et al. (2008) | 10.1097/WNR.0b013e3282ffda89 | 21 | visual | complexity | MNI | center-embedded > left-branching | -36 | 10 | 40 |
|  |  |  |  |  |  |  | -40 | 18 | 28 |
|  |  |  |  |  |  | center embedded > active-co-joined sentence | -34 | 8 | 44 |
|  |  |  |  |  |  |  | -28 | -52 | 42 |
|  |  |  |  |  |  |  | 32 | -48 | 50 |
|  |  |  |  |  |  |  | 30 | 6 | 50 |
|  |  |  |  |  |  |  | 0 | 26 | 54 |
|  |  |  |  |  |  |  | 10 | -74 | -36 |
|  |  |  |  |  |  |  | 38 | 16 | 28 |
| Ohta 2017 | 10.3389/fpsyg.2017.00748 | 17 | visual & auditory | complexity | MNI | VSO > VOS | -36 | 6 | 57 |
|  |  |  |  |  |  |  | -48 | 0 | 45 |
|  |  |  |  |  |  |  | -48 | 12 | 42 |
|  |  |  |  |  |  |  | -57 | 12 | 12 |
|  |  |  |  |  |  |  | -54 | 24 | 0 |
|  |  |  |  |  |  |  | -6 | 15 | 48 |
|  |  |  |  |  |  |  | -12 | -78 | 45 |
|  |  |  |  |  |  |  | -27 | -69 | 39 |
|  |  |  |  |  |  |  | -30 | -75 | 24 |
|  |  |  |  |  |  |  | 18 | -69 | 51 |
|  |  |  |  |  |  |  | 30 | -66 | 51 |
|  |  |  |  |  |  | OVS > SVO | -42 | 3 | 54 |
|  |  |  |  |  |  |  | -51 | 18 | 27 |
|  |  |  |  |  |  |  | -54 | 30 | 0 |
|  |  |  |  |  |  |  | -3 | 15 | 60 |
|  |  |  |  |  |  |  | -6 | 15 | 48 |
|  |  |  |  |  |  |  | -3 | 27 | 42 |
|  |  |  |  |  |  | VSO + OVS > VOS + SVO | -42 | 3 | 51 |
|  |  |  |  |  |  |  | -45 | 12 | 42 |
|  |  |  |  |  |  |  | -36 | 12 | 33 |
|  |  |  |  |  |  |  | -48 | 18 | 27 |
|  |  |  |  |  |  |  | -54 | 12 | 15 |
|  |  |  |  |  |  |  | -54 | 27 | 0 |
|  |  |  |  |  |  |  | -6 | 15 | 48 |
|  |  |  |  |  |  |  | 0 | 24 | 42 |
|  |  |  |  |  |  |  | -27 | -69 | 42 |
| Pattamadilok 2016 | 10.1016/j.cortex.2015.11.012 | 20 | visual | complexity | MNI | embedded > adjunct sentence during probe presentation | -48 | 20 | 28 |
|  |  |  |  |  |  |  | -54 | 20 | 19 |
|  |  |  |  |  |  |  | -48 | 29 | 22 |
|  |  |  |  |  |  |  | -57 | 20 | 1 |
|  |  |  |  |  |  |  | 54 | 26 | 28 |
|  |  |  |  |  |  |  | -48 | 26 | -8 |
|  |  |  |  |  |  |  | -48 | 38 | -8 |
|  |  |  |  |  |  |  | -27 | 23 | -5 |
|  |  |  |  |  |  |  | 33 | 26 | -5 |
|  |  |  |  |  |  |  | -57 | -37 | -2 |
|  |  |  |  |  |  |  | -51 | -46 | 4 |
|  |  |  |  |  |  | embedded > adjunct sentence during related probe presentation | -27 | 23 | -2 |
|  |  |  |  |  |  |  | -48 | 38 | -8 |
|  |  |  |  |  |  |  | -54 | 23 | 19 |
|  |  |  |  |  |  |  | -51 | 17 | 10 |
|  |  |  |  |  |  |  | -51 | 14 | 40 |
|  |  |  |  |  |  |  | 33 | 23 | -5 |
|  |  |  |  |  |  |  | 54 | 23 | 34 |
|  |  |  |  |  |  |  | -39 | 53 | 1 |
|  |  |  |  |  |  |  | 0 | 32 | 49 |
|  |  |  |  |  |  |  | -3 | 14 | 55 |
|  |  |  |  |  |  |  | -3 | 5 | 67 |
|  |  |  |  |  |  |  | -39 | 2 | 43 |
|  |  |  |  |  |  |  | -57 | -43 | 1 |
|  |  |  |  |  |  |  | -36 | -55 | 46 |
|  |  |  |  |  |  |  | 3 | -64 | 46 |
| Peelle 2004 | 10.1016/j.bandl.2004.05.007 | 8 | auditory | complexity | TAL | Object-relative > Subject-relative center-embedded clause | -44 | 15 | -7 |
|  |  |  |  |  |  |  | 16 | 23 | -8 |
| Prat et al. (2011) | 10.1093/cercor/bhq241 | 27 | visual | complexity | MNI | Sentence with object-relative clause > co-joined clauses | -44 | 16 | 14 |
|  |  |  |  |  |  |  | -26 | 30 | -2 |
|  |  |  |  |  |  |  | -40 | 4 | 48 |
|  |  |  |  |  |  |  | -6 | 10 | 64 |
|  |  |  |  |  |  |  | -12 | 58 | 24 |
|  |  |  |  |  |  |  | 6 | 50 | -16 |
|  |  |  |  |  |  |  | -58 | -10 | -20 |
|  |  |  |  |  |  |  | 36 | 26 | 12 |
|  |  |  |  |  |  |  | 54 | 30 | 0 |
|  |  |  |  |  |  |  | 34 | -2 | 44 |
|  |  |  |  |  |  |  | 50 | -10 | -26 |
|  |  |  |  |  |  |  | 50 | -46 | 10 |
|  |  |  |  |  |  |  | 26 | -32 | 26 |
|  |  |  |  |  |  |  | -12 | -12 | -18 |
|  |  |  |  |  |  |  | 22 | -6 | -14 |
|  |  |  |  |  |  |  | 8 | 12 | 14 |
|  |  |  |  |  |  |  | 8 | -2 | -8 |
|  |  |  |  |  |  |  | -30 | -32 | 14 |
|  |  |  |  |  |  |  | -10 | -60 | -28 |
| Quinones 2014 | 10.1016/j.neuroimage.2013.11.038 | 21 | visual | violation | MNI | person mismatch > unagreement | 2 | 34 | 34 |
|  |  |  |  |  |  |  | -44 | 22 | 36 |
|  |  |  |  |  |  |  | -40 | 46 | 22 |
|  |  |  |  |  |  |  | -32 | 46 | 34 |
|  |  |  |  |  |  |  | -42 | -46 | 58 |
|  |  |  |  |  |  |  | -6 | -66 | 50 |
|  |  |  |  |  |  |  | 10 | -32 | 52 |
|  |  |  |  |  |  |  | 6 | 40 | 26 |
|  |  |  |  |  |  |  | 56 | -42 | 50 |
|  |  |  |  |  |  |  | 58 | -36 | 44 |
| Quinones 2018 | 10.1016/j.neuroimage.2018.03.069 | 47 | visual | violation | MNI | gender mismatch > match | -4 | 52 | -2 |
|  |  |  |  |  |  |  | -26 | 24 | 50 |
|  |  |  |  |  |  |  | -6 | 38 | -6 |
|  |  |  |  |  |  |  | -6 | -22 | 60 |
|  |  |  |  |  |  |  | -42 | -6 | 32 |
|  |  |  |  |  |  |  | -44 | -16 | 34 |
|  |  |  |  |  |  |  | -48 | -66 | 42 |
|  |  |  |  |  |  |  | -4 | -48 | 10 |
|  |  |  |  |  |  |  | -8 | -40 | 26 |
|  |  |  |  |  |  |  | -16 | -82 | 28 |
|  |  |  |  |  |  |  | -4 | -74 | -2 |
|  |  |  |  |  |  |  | 10 | 52 | 2 |
|  |  |  |  |  |  |  | 26 | 54 | 6 |
|  |  |  |  |  |  |  | 50 | 12 | 42 |
|  |  |  |  |  |  |  | 34 | -2 | 16 |
|  |  |  |  |  |  |  | 14 | 14 | 12 |
|  |  |  |  |  |  |  | 26 | 8 | 10 |
|  |  |  |  |  |  |  | 2 | -16 | 68 |
|  |  |  |  |  |  |  | 8 | -70 | -4 |
| Raettig 2010 | 10.1016/j.cortex.2009.06.003 | 15 | auditory | violation | TAL | morphosyntactically incorrect > correct | -65 | -42 | 15 |
| Rogalsky 2008 | 10.3389/neuro.09.014.2008 | 15 | visual | complexity | TAL | object-relative > subject-relative sentence | -41 | 38 | 14 |
|  |  |  |  |  |  |  | -42 | 13 | 23 |
|  |  |  |  |  |  |  | -56 | -37 | 19 |
|  |  |  |  |  |  |  | -54 | 4 | 31 |
|  |  |  |  |  |  |  | -39 | 32 | 32 |
|  |  |  |  |  |  |  | -32 | 56 | 19 |
|  |  |  |  |  |  |  | 47 | 16 | 20 |
| Röder et al. (2002) | 10.1006/nimg.2001.1026 | 11 | auditory | complexity | TAL | syntactically difficult > easy (word order) | -45 | 12 | 16 |
|  |  |  |  |  |  |  | -47 | -45 | 9 |
|  |  |  |  |  |  |  | -44 | 3 | 36 |
|  |  |  |  |  |  |  | -2 | 6 | 50 |
|  |  |  |  |  |  |  | 31 | 19 | 2 |
| Seyfried 2023 | 10.1080/23273798.2022.2116462 | 25 | auditory | violation | MNI | Syntactic violation > correct | -32 | 28 | -2 |
|  |  |  |  |  |  |  | -38 | 22 | 0 |
|  |  |  |  |  |  |  | -46 | 8 | 6 |
|  |  |  |  |  |  |  | -16 | 32 | 28 |
|  |  |  |  |  |  |  | -6 | 10 | 56 |
|  |  |  |  |  |  |  | -8 | 14 | 28 |
|  |  |  |  |  |  |  | 14 | -30 | 26 |
|  |  |  |  |  |  |  | -6 | -30 | 24 |
|  |  |  |  |  |  |  | 22 | -34 | 30 |
| Shetreet 2009 | 10.1016/j.neuroimage.2009.07.001 | 19 | auditory | complexity | TAL | sentence with sentencial complement > sentence with noun phrase complement | -56 | 21 | 7 |
|  |  |  |  |  |  |  | -48 | -35 | 5 |
|  |  |  |  |  |  |  | 45 | -24 | -6 |
|  |  |  |  |  |  |  | -53 | -15 | -14 |
|  |  |  |  |  |  |  | 50 | 2 | -25 |
|  |  |  |  |  |  |  | -59 | -51 | 27 |
|  |  |  |  |  |  |  | 56 | -48 | 22 |
|  |  |  |  |  |  |  | -9 | -54 | 33 |
|  |  |  |  |  |  |  | -12 | -25 | 29 |
|  |  |  |  |  |  |  | -18 | -9 | 0 |
|  |  |  |  |  |  |  | -18 | 45 | 34 |
|  |  |  |  |  |  |  | -30 | 5 | 38 |
| Shetreet 2014 | 10.1016/j.jneuroling.2013.06.003 | 22 | auditory | complexity | TAL | wh-movement > canonical order | -53 | 24 | 10 |
|  |  |  |  |  |  |  | 50 | 27 | -5 |
|  |  |  |  |  |  |  | -53 | -27 | 0 |
|  |  |  |  |  |  |  | 51 | -35 | 2 |
|  |  |  |  |  |  |  | -12 | -81 | -26 |
|  |  |  |  |  |  |  | -48 | -1 | 50 |
|  |  |  |  |  |  | verb movement > canonical order | -18 | -94 | -7 |
| Snijders 2009 | 10.1093/cercor/bhn187 | 22 | visual | ambiguity | MNI | word-class ambiguous > unambiguous sentence | -52 | -50 | -8 |
|  |  |  |  |  |  |  | -60 | -44 | -4 |
|  |  |  |  |  |  |  | -46 | -54 | -4 |
|  |  |  |  |  |  |  | -50 | -20 | -28 |
|  |  |  |  |  |  |  | -42 | -24 | -28 |
|  |  |  |  |  |  |  | -46 | -30 | -22 |
|  |  |  |  |  |  |  | 48 | -34 | -14 |
|  |  |  |  |  |  |  | 50 | -44 | -4 |
|  |  |  |  |  |  |  | -44 | 0 | 22 |
|  |  |  |  |  |  |  | 46 | 28 | 6 |
|  |  |  |  |  |  |  | 44 | 18 | 14 |
|  |  |  |  |  |  |  | 54 | 18 | 12 |
| Stowe 2004 | 10.1016/S0093-934X(03)00359-6 | 16 | visual | ambiguity | TAL | syntactically ambiguous > unambiguous sentences | -48 | 20 | 28 |
|  |  |  |  |  |  |  | 18 | 6 | 16 |
|  |  |  |  |  |  |  | 38 | -80 | -24 |
|  |  |  |  |  |  |  | -14 | 36 | 44 |
| Stromswold 1996 | 10.1006/brln.1996.0024 | 8 | visual | complexity | TAL | center-embedded > right-branching sentence | 46.50 | 9.80 | 4.00 |
| Suh et al. (2007) | 10.1016/j.brainres.2006.12.043 | 16 | visual | complexity | TAL | embedded > conjoined sentence | -38 | -52 | 46 |
|  |  |  |  |  |  |  | -8 | -70 | 46 |
|  |  |  |  |  |  |  | -46 | 16 | 46 |
| Tanaka 2017 | 10.2183/pjab.93.031 | 16 | visual | complexity | MNI | object scrambling > unscrambling sentence | -42 | -1 | 26 |
|  |  |  |  |  |  |  | -48 | 14 | 20 |
|  |  |  |  |  |  |  | -60 | 11 | 17 |
| Thibault 2021 | 10.1126/science.abe0874 | 20 | visual | complexity | MNI | 2 object-relative clauses > (coordinated clauses + subject-relative clauses | -18 | 14 | -1 |
|  |  |  |  |  |  |  | -42 | 29 | 5 |
|  |  |  |  |  |  |  | -42 | -52 | 32 |
|  |  |  |  |  |  |  | 15 | 11 | -1 |
|  |  |  |  |  |  |  | 51 | -49 | 26 |
| Xiong 2021 | 10.1016/j.neuroimage.2020.117475 | 29 | visual | complexity | MNI | center-embedding object relative clause > left-branching object relative clause | -42 | -60 | 48 |
|  |  |  |  |  |  |  | -38 | 14 | 26 |
|  |  |  |  |  |  |  | 46 | -40 | 42 |
|  |  |  |  |  |  | center-embedding object relative clause > center-embedding subject-relative clause | -56 | -8 | -14 |
|  |  |  |  |  |  |  | -48 | -58 | 22 |
| Xu 2020 | 10.1016/j.bandl.2019.104712 | 19 | visual | complexity | MNI | hard subject-extracted relative clause - visual baseline > easy object-extracted relative clause - visual baseline in Chinese | -34 | 28 | -4 |
|  |  |  |  |  |  |  | -60 | -50 | 16 |
| Ye and Zhou (2009) | 10.1016/j.neuroimage.2009.06.032 | 19 | visual | complexity | MNI | Passive > active sentences | -12 | 8 | 60 |
|  |  |  |  |  |  |  | -54 | 22 | 14 |
|  |  |  |  |  |  |  | -32 | 30 | -4 |
|  |  |  |  |  |  |  | 34 | -88 | -10 |
| Zhang 2024 | 10.1016/j.neuroimage.2024.120543 | 29 | visual | complexity | MNI | marked non-canonical > canonical word order sentence | -54 | -42 | 0 |
|  |  |  |  |  |  |  | -42 | 0 | 45 |
|  |  |  |  |  |  |  | -54 | 18 | 18 |
|  |  |  |  |  |  |  | -42 | 27 | -9 |
|  |  |  |  |  |  |  | -3 | 15 | 51 |
|  |  |  |  |  |  | unmarked non-canonical with same-animacy arguments > canonical sentence | -54 | 15 | 18 |
|  |  |  |  |  |  |  | -39 | 3 | 39 |
|  |  |  |  |  |  |  | -54 | 15 | 54 |
|  |  |  |  |  |  |  | -54 | 18 | 6 |
|  |  |  |  |  |  |  | -51 | 33 | -9 |
|  |  |  |  |  |  |  | -48 | 9 | -9 |
|  |  |  |  |  |  |  | -36 | -48 | 36 |
|  |  |  |  |  |  |  | -42 | 0 | 45 |
|  |  |  |  |  |  |  | -54 | -48 | 0 |
|  |  |  |  |  |  |  | -63 | -51 | 45 |
|  |  |  |  |  |  |  | -48 | 18 | 48 |
|  |  |  |  |  |  |  | 30 | -66 | -54 |
|  |  |  |  |  |  |  | -30 | 3 | -33 |
|  |  |  |  |  |  |  | 42 | 3 | 33 |
|  |  |  |  |  |  | unmarked non-canonical with different animacy argument > canonical sentence | -54 | -39 | -3 |
|  |  |  |  |  |  |  | -36 | -48 | -3 |
|  |  |  |  |  |  |  | -48 | -42 | 36 |
|  |  |  |  |  |  |  | -54 | 18 | -9 |
|  |  |  |  |  |  |  | -51 | 30 | -6 |
|  |  |  |  |  |  |  | -27 | 0 | 48 |
|  |  |  |  |  |  |  | -39 | 0 | 33 |
|  |  |  |  |  |  |  | -54 | 18 | 15 |
|  |  |  |  |  |  |  | -6 | 12 | 57 |

Note: TAL = talairach space.

**Syntax violation and complexity**

To assess whether this result was driven by a single type of task contrast or true across the different assessments of syntax, we compared the two main contrast types: syntactic complexity and violation (Supplementary Figure 1 and Table 2). The two studies on syntactic ambiguity were excluded for this step only. Assessments of syntactic complexity recruits a very similar network to the overall syntax finding, including left IFG (pars opercularis and pars triangularis) extending into precentral gyrus and MFG, pSTS/MTG, dmPFC, and IPL, as well as right IFG (pars orbitalis)/insula. This is not surprising as this includes the majority of the syntax studies (50 out of 66). In contrast, the syntactic violation network is quite limited due to it only including 14 studies. The two contrast types overlap in a small cluster in the left IFG (pars opercularis). While there is significantly greater involvement of a small area of left IFG (pars triangularis) for the complexity studies this could be due to the lack of power in the violation assessment.

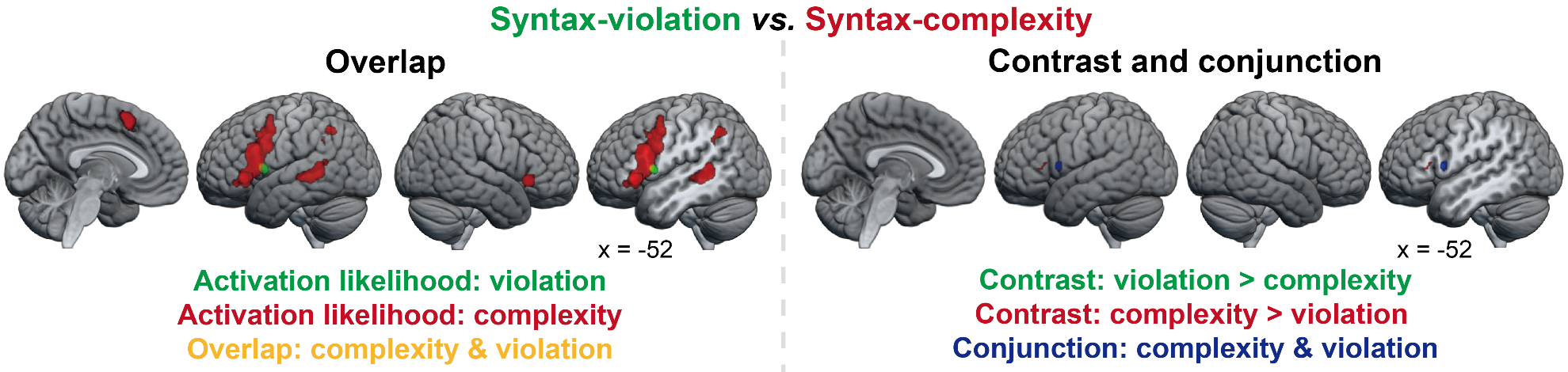

**Supplementary Figure 1** Overlap of ALE results for violation and complexity (left, overlap in yellow). Contrast and conjunction analysis between violation and complexity at a voxel-level *p* < 0.001 with 10,000 permutations, cluster volume > 20 mm^3^ right, auditory violation > complexity in green, complexity> violation in red, conjunction in blue).

**Supplementary Table 2** Contrast analyses between syntax complexity and syntax violation

| Cluster number | Region | Peak MNI Coordinates | | |
| --- | --- | --- | --- | --- |
|  |  | X | Y | Z |
| *Contrast analysis: Syntax complexity > syntax violation* | | | | |
| 1 | L IFG (pars triangularis) | -53 | 30 | 6 |
|  |  | -51 | 24 | 10 |
| *Contrast analysis: Syntax violation > syntax complexity* | | | | |
| No significant results | | | | |
| *Conjunction analysis: syntax violation & syntax complexity* | | | | |
| 1 | L IFG (pars opercularis) | -50 | 10 | 8 |

Note: IFG = inferior frontal gyrus. Voxel-level *p* < 0.001 with 10,000 permutations, cluster volume > 20 mm^3^.
